# Human parietal retrieval states prioritize an absent temporal reference to reconstruct event order

**DOI:** 10.64898/2026.08.11.744125

**Authors:** Chengmei Huang, Jing Fu, Ruimin Wang, Isao Hasegawa, Koji Jimura, Kiyoshi Nakahara

## Abstract

How the brain judges when something happened is thought to involve reconstruction—deriving temporal judgments from information recovered from memory rather than simply reading out a fixed temporal code. Testing this has been difficult because conventional tasks present both compared items at retrieval, so reinstatement could reflect cue processing as much as memory recovery. Here we dissociated what a judgment requires from what its cue supplies: during fMRI scanning, participants (n = 55) judged a target’s temporal position relative to a milestone that was never shown at retrieval. We found that retrieval patterns in the precuneus and angular gyrus resembled the absent milestone more than the target, a bias strongest in the precuneus. Moreover, trial-by-trial similarity to the encoded path linking milestone to target—rather than to either alone—predicted correct judgments independently of retrieval activation. Human parietal retrieval is thus configured more by what a judgment requires than by what the cue supplies, providing content-specific representational evidence for reconstructive temporal-order memory.

## Introduction

Episodic memory is defined not only by what happened, but by when it happened and how events followed one another in time ^1,2^. Yet memories do not appear to carry a single, explicit timestamp. Reconstructive accounts propose that judgments about when past events occurred are inferred from multiple sources of information available in memory, rather than read out from a single, fixed temporal code ^3,4^. Consistent with this reconstructive view, temporal-order memory is shaped by the contextual organization of event sequences ^5^, is associated with encoding-stage temporal-context representations ^6^, and can involve reactivation of information tied to items intervening between the two judged events ^7^.

In representational terms, reconstruction fundamentally involves the active recovery of temporally useful mnemonic information at retrieval. Testing this has proven difficult, for a reason built into the standard TOJ paradigm. In conventional TOJ tasks, the two items whose temporal relation must be judged are typically displayed as retrieval cues ^6–9^. For the judged items themselves, the information that a reconstructive account predicts should be internally recovered is precisely the information the display supplies; apparent reinstatement may therefore reflect perceptual processing of the cues rather than, or in addition to, mnemonic recovery ^10^. A related ambiguity applies to information that is not itself displayed. When such information is reinstated ^7^, it remains unclear whether it was selectively recovered for the current judgment or passively carried along by cue-triggered associative or temporal-context retrieval ^11–13^. The reconstructive account has therefore remained effectively untested at the representational level.

We resolved this ambiguity by inverting the relationship between what is shown and what is needed. In the reference-based TOJ task introduced here, participants encoded sequences of nine items, each containing a milestone at the central serial position that served as a task-defined temporal reference point. At retrieval, only the name of a single target object was presented, and participants judged the target’s encoded temporal position relative to the milestone. The judgment therefore could not be made from the cue alone: the anchor against which temporal position had to be evaluated was absent from the display and available only from memory. If retrieval states are configured by the current mnemonic goal rather than by perceptual input, they should be more closely aligned with the absent reference point than with the target named by the on-screen cue. Such a pattern could not be attributed to direct presentation of the reference point and would indicate that retrieval was not dominated by the externally presented target.

Posterior parietal cortex is a natural place to look for such goal-configured retrieval states. The precuneus (PCu) is consistently engaged during episodic retrieval ^14–17^, and its activity varies with the temporal distance between remembered events ^18^, reflects the temporal proximity and relevance of information encountered during retrieval ^19^, and tracks storyline context ^20^. PCu activity is also greater for short-lag autobiographical temporal-order judgments, which has been interpreted as reflecting greater demands on reconstructive retrieval ^21^. Recent macaque work has further shown that population-level encoding–retrieval similarity in medial posterior parietal cortex (mPPC; the precuneus) predicts temporal-order judgment performance ^22^, providing convergent cross-species evidence for a medial parietal contribution to temporal-order retrieval. The neighboring angular gyrus (AG) has likewise been linked to successful recollection ^23^ and to multimodal feature integration during episodic retrieval ^24^, as well as to temporal distance and narrative context ^18,20^. What this body of work leaves open is not whether posterior parietal retrieval states reinstate encoding-related information, but whether that information reflects the demands of the current judgment rather than the perceptual or associative pull of the retrieval cue—the question on which the reconstructive account turns. The Attention-to-Memory (AtoM) model anticipates this expectation ^25,26^: if parietal cortex orients mnemonic processing toward goal-relevant content, retrieval states should be shaped by what the judgment demands rather than by what the cue provides.

The present fMRI study tested two hierarchically ordered predictions derived from reconstructive accounts using encoding–retrieval similarity (ERS) analysis ^27,28^ in a priori bilateral PCu and AG regions of interest, complemented by whole-brain searchlights. First, if retrieval is organized around the internally required anchor, retrieval patterns should preferentially reinstate the encoding pattern of the absent reference point—over the other items of the same sequence and, critically, over the externally presented target. Second, and more stringently, recovering the anchor alone cannot specify where the target stood: reconstruction requires the encoded material that links anchor to target. Trial-wise similarity to the encoding patterns of the reference-to-target sequence segment, rather than to either endpoint alone or to the remainder of the sequence, should therefore track judgment accuracy. Both predictions were borne out. Retrieval patterns in PCu and AG were biased toward the absent reference point over both the other sequence items and the presented target, with the bias stronger in PCu, and path-specific similarity predicted correct judgments in both regions. These results provide representational evidence for a reconstructive account of temporal-order retrieval, in which posterior parietal states are configured by the structure of the judgment itself rather than simply following the information specified by the external cue.

## Results

### Behavioral performance varied by distance from the milestone

Participants completed a reference-based TOJ task comprising encoding, interference, and retrieval phases (Fig. 1). During encoding, they studied sequences of eight object images and one milestone image. The milestone always appeared at the middle serial position and served as the temporal reference point. During retrieval, the name of one object from the just-encoded sequence cued participants to report the target’s position relative to the milestone. Using object names as retrieval cues reduced perceptual overlap between encoding and retrieval, and each encoded sequence was tested once to minimize retrieval-practice effects and carryover across retrieval episodes. Trials were classified according to the target’s relative position (before vs. after the milestone) and temporal distance from the milestone: D1, D2, and D3 denoted targets immediately adjacent to the milestone, separated by one intervening object, or separated by two intervening objects, respectively (see Methods for details).

**Fig. 1.**
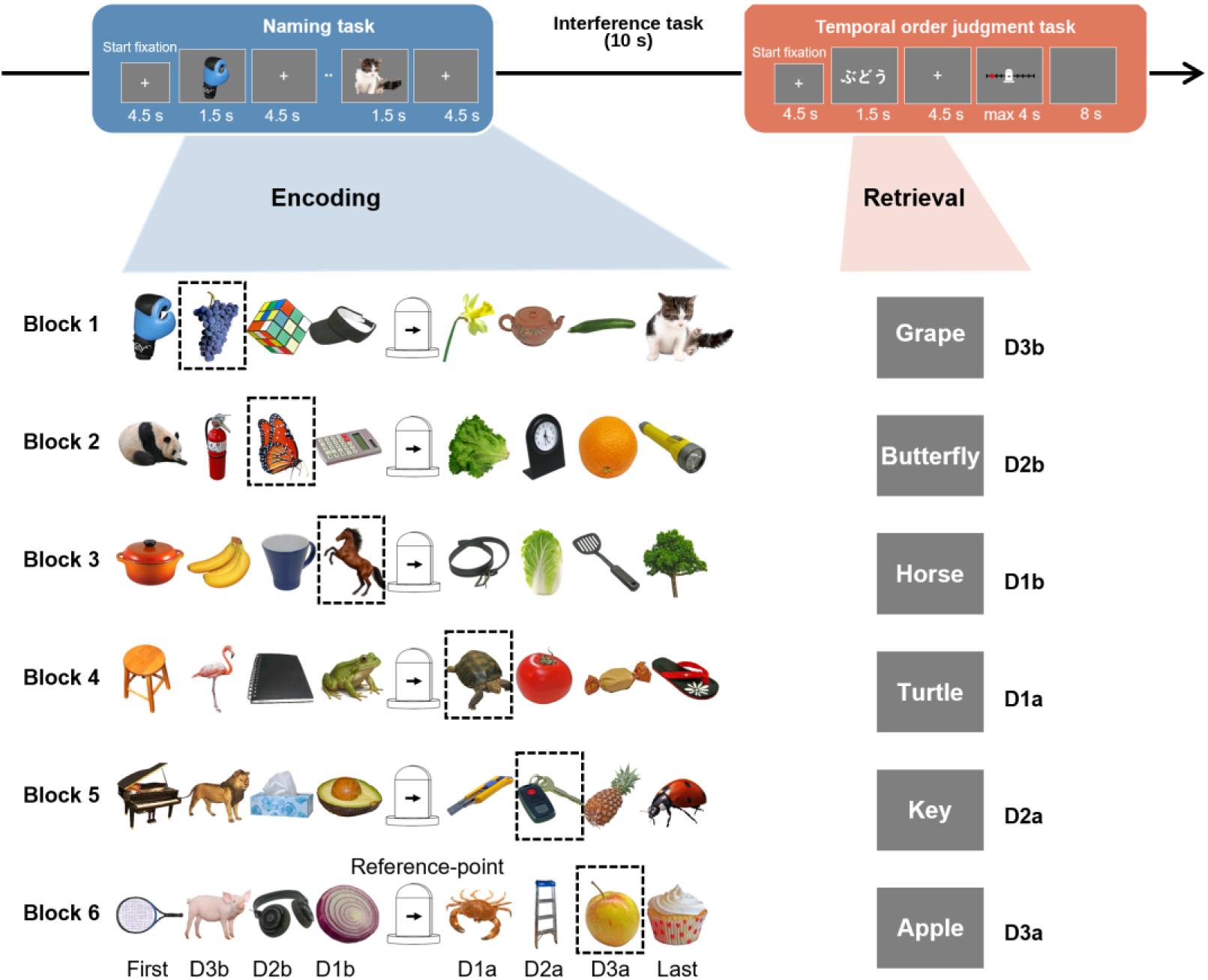
Reference-based temporal-order judgment task. The experiment comprised four runs of six blocks, each with encoding, interference and retrieval phases. During encoding, participants studied a nine-item sequence containing eight objects and a white gate milestone at serial position 5, which served as the temporal reference point. During retrieval, the name of one target object cued participants to report its encoded position relative to the absent milestone on a response timeline. Trial types were defined by temporal distance and direction relative to the milestone: D1, D2 and D3 denote zero, one and two intervening objects, respectively, and b and a denote before and after. Dashed outlines identify the target in each example and were not shown during the task. English object names are shown for illustration; retrieval cues were presented in Japanese.

We analyzed TOJ accuracy during fMRI scanning as a function of relative position and temporal distance. A 2 × 3 repeated-measures ANOVA with relative position and temporal distance as within-participant factors showed a significant main effect of temporal distance (F(1.98, 106.68) = 6.18, p = 0.003, ηG² = 0.022), no significant main effect of relative position (F(1, 54) = 1.82, p = 0.183, ηG² = 0.005), and a significant relative position × temporal distance interaction (F(1.99, 107.84) = 7.88, p < 0.001, ηG² = 0.041; Fig. 2a). Bonferroni-corrected post hoc comparisons showed that this interaction was driven by after-milestone targets: accuracy was lower for D3a than for D1a (t(54) = 5.25, p_Bonf_ < 0.001, Hedges’ g = 0.85) and D2a (t(54) = 4.23, p_Bonf_ < 0.001, Hedges’ g = 0.65), whereas D1a and D2a did not differ (t(54) = 1.08, p = 0.286, Hedges’ g = 0.17). No significant differences were observed among before-milestone targets (all p > 0.3). Thus, accuracy differed by distance only for after-milestone targets, with poorer performance for D3a than for targets closer to the milestone.

**Fig. 2.**
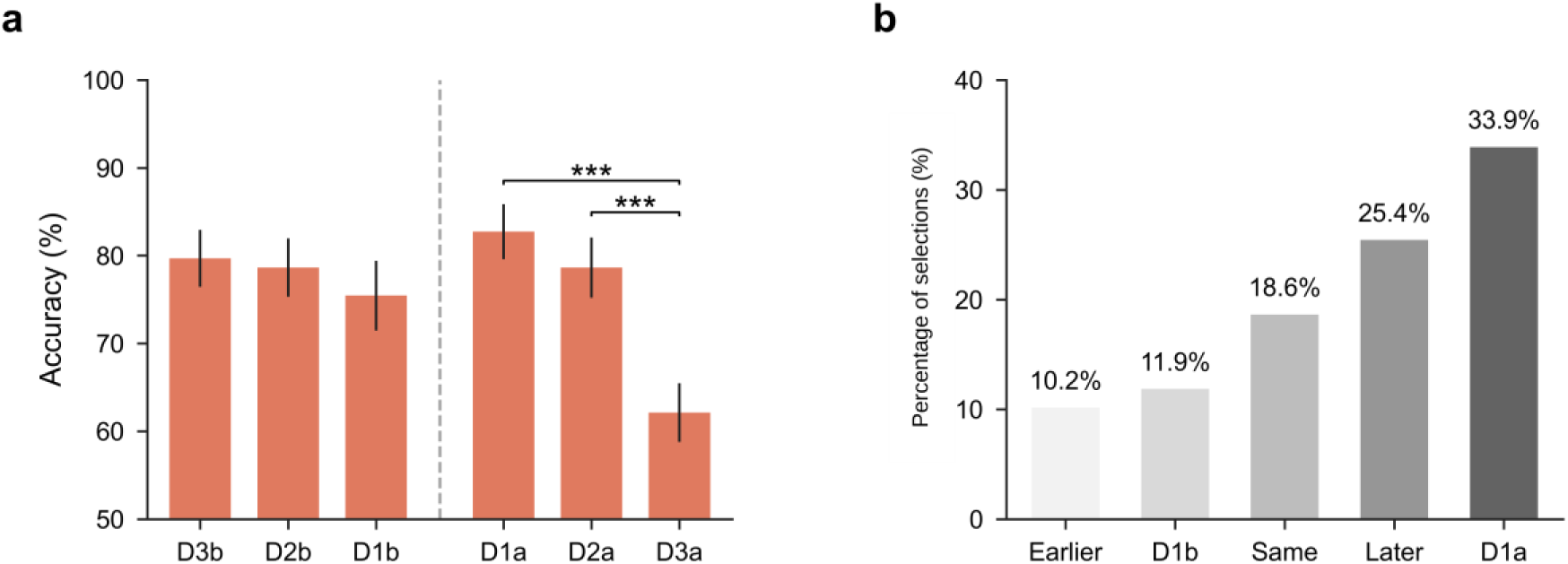
Behavioral performance in the reference-based temporal-order judgment task. **a,** Mean temporal-order judgment (TOJ) accuracy across the six trial types, ordered by target position relative to the milestone. The dashed vertical line separates before- and after-milestone targets. Error bars show s.e.m. across participants (*n* = 55). Brackets denote Bonferroni-corrected two-sided post hoc comparisons (***p_Bonf_ < 0.001). **b,** Distribution of post-scan reports of the easiest temporal position. Multiple responses were permitted, yielding 59 selections from 55 participants. Earlier and Later indicate positions earlier or later in the sequence; D1b and D1a indicate positions immediately before and after the milestone; Same indicates no perceived positional difference. Values above bars show percentages of all selections.

After scanning, participants completed an exploratory questionnaire about which target position was easiest to remember (Fig. 2b). Multiple responses were permitted, yielding 59 selections from 55 participants. The distribution of selections was not uniform across target positions (chi-square-type statistic = 11.42, permutation p = 0.019). Follow-up tests showed that D1a was selected 20 times (33.9% of all selections), more often than expected under the null model (permutation p = 0.0078, p_Bonf_ = 0.039), whereas no other option survived Bonferroni correction (all p_Bonf_ > 0.90). Both accuracy and questionnaire responses were sensitive to the milestone-based temporal structure, with the clearest advantage near the milestone for after-milestone targets.

### Univariate results

Before testing the two ERS predictions, we examined univariate activation in four a priori posterior parietal ROIs: left PCu, right PCu, left AG, and right AG (Fig. 3a). These analyses tested whether PCu and AG were engaged during temporal-order retrieval and whether retrieval activation in these regions was related to TOJ performance.

**Fig. 3.**
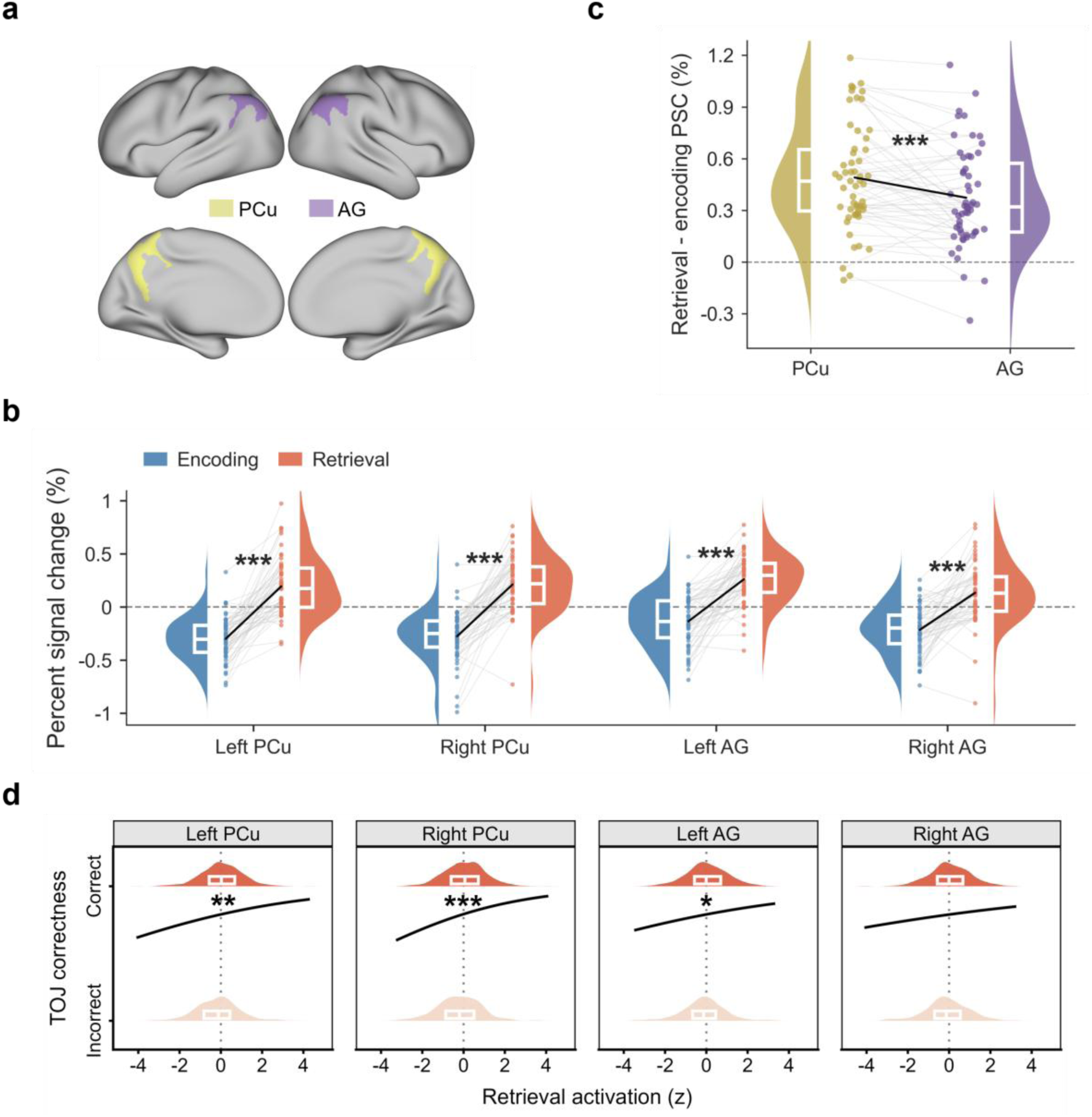
Posterior parietal activation during temporal-order retrieval. **a,** A priori bilateral precuneus (PCu, yellow) and angular gyrus (AG, purple) regions of interest (ROIs) shown on standard cortical surfaces; analyses used participant-native Destrieux atlas masks. *n* = 55 participants. **b,** Percent signal change (PSC) during encoding and retrieval in each ROI. Half-violin plots and boxplots show participant-level distributions, gray lines connect paired observations and black lines connect group means. Asterisks denote FDR-corrected two-sided paired *t* tests (*p_FDR_ < 0.05, **p_FDR_ < 0.01, ***p_FDR_ < 0.001). **c,** Retrieval-minus-encoding PSC, averaged across hemispheres, in PCu and AG. Half-violin plots and boxplots show participant-level distributions, gray lines connect paired observations and black lines connect group means. ***p < 0.001, two-sided PCu–AG comparison. **d,** Trial-wise association between retrieval activation and temporal-order judgment (TOJ) correctness. Marginal density plots and boxplots show retrieval-activation distributions for incorrect and correct trials; black curves show population-level model-predicted probabilities of a correct TOJ, adjusted for trial type. Boxplots show medians and interquartile ranges. Asterisks denote FDR-corrected fixed effects from mixed-effects logistic regression models (*p_FDR_ < 0.05, **p_FDR_ < 0.01, ***p_FDR_ < 0.001).

### Temporal-order retrieval engaged bilateral PCu and AG, with stronger activation increases in PCu

Percent signal change (PSC) was extracted from each ROI for the matched encoding-phase target-object and retrieval-phase cue regressors. All four ROIs showed greater activation during retrieval than encoding (Fig. 3b): left PCu (t(54) = 10.60, p_FDR_ = 2.132 × 10^−14^, Hedges’ g = 1.42), right PCu (t(54) = 10.99, p_FDR_ = 1.110 × 10^−14^, Hedges’ g = 1.48), left AG (t(54) = 9.48, p_FDR_ = 7.049 × 10^−13^, Hedges’ g = 1.27), and right AG (t(54) = 7.59, p_FDR_ = 5.152 × 10^−10^, Hedges’ g = 1.02).

We then tested whether this retrieval activation increase was larger in PCu than in AG. A 2 × 2 repeated-measures ANOVA with brain region (PCu vs. AG) and hemisphere (left vs. right) as within-participant factors showed a significant main effect of brain region (F(1, 54) = 13.30, p < 0.001, ηG² = 0.032), reflecting a larger activation increase in PCu than in AG (paired comparison: t(54) = 3.65, p < 0.001, Hedges’ g = 0.39). Neither the main effect of hemisphere nor the brain region × hemisphere interaction was significant (both p > 0.2; Fig. 3c).

As a complementary whole-brain analysis of univariate activation, we tested whether the retrieval > encoding effect observed in the predefined ROIs was also evident beyond these regions. The contrast revealed a distributed network spanning bilateral posterior parietal, frontal, occipitotemporal, cingulate, sensorimotor, subcortical and cerebellar regions, including clusters overlapping the predefined PCu and AG regions (p_FWE_ < 0.05, TFCE-corrected; Supplementary Fig. 1 and Supplementary Table 1). This distribution partially overlapped with regions previously implicated in temporal retrieval and cue-guided reconstruction of event structure ^18,20,29^.

### Retrieval activation was associated with TOJ correctness, especially in PCu

We next tested whether trial-wise cue-evoked activation during the retrieval phase was associated with TOJ correctness (correct = 1, incorrect = 0). For comparison, we tested whether trial-wise activation evoked by the target object during the encoding phase was similarly associated with TOJ correctness. Separate mixed-effects logistic regression models were fitted for each neural measure and predefined ROI, with trial type (D3b, D2b, D1b, D1a, D2a, and D3a) included as a fixed effect and a participant-specific random intercept to account for repeated observations within participants (see Methods for details).

Retrieval activation was positively associated with trial-wise TOJ correctness in left PCu (z = 3.00, p_FDR_ = 0.005; odds ratio = 1.24, 95% CI = [1.08, 1.43]), right PCu (z = 4.10, p_FDR_ < 0.001; odds ratio = 1.35, 95% CI = [1.17, 1.55]), and left AG (z = 2.51, p_FDR_ = 0.016; odds ratio = 1.20, 95% CI = [1.04, 1.38]; Fig. 3d; Supplementary Table 2). Right AG showed a weaker association that did not survive FDR correction (z = 1.83, p_FDR_ = 0.068; odds ratio = 1.14, 95% CI = [0.99, 1.31]). Encoding activation was not significantly associated with trial-wise TOJ correctness in any ROI (all p_FDR_ ≥ 0.237; Supplementary Table 3).

To assess whether PCu and AG retrieval activation showed independent associations with trial-wise TOJ correctness, we fitted a mixed-effects logistic regression model that included bilateral PCu retrieval activation and bilateral AG retrieval activation as simultaneous predictors. Variance inflation factors were below 2 for both predictors, indicating low collinearity. After mutual adjustment, bilateral PCu retrieval activation remained positively associated with trial-wise TOJ correctness (z = 2.89, odds ratio = 1.34, 95% CI = [1.10, 1.64], p = 0.004), whereas bilateral AG retrieval activation showed no independent association (z = −0.41, p = 0.680).

As a complementary whole-brain searchlight analysis, we tested whether the ROI-level association between cue-evoked retrieval activation and trial-wise TOJ correctness extended beyond the predefined ROIs. Positive associations were observed in a bilateral PCu/superior parietal cluster, right angular/inferior parietal and angular/middle occipital regions, along with additional frontal/precentral and left cerebellar clusters. Negative associations were observed in left middle occipital/middle temporal cortex, left caudate, and right precentral/middle frontal cortex (FDR q < 0.05, cluster size ≥ 20 voxels; see Supplementary Fig. 2 and Supplementary Table 4 for details).

### Multivariate results

The preceding univariate analyses showed that bilateral PCu and AG were engaged during temporal-order retrieval and that greater retrieval activation was associated with correct TOJ, most robustly in PCu. These analyses established retrieval engagement but could not identify which encoding-related information was expressed in the retrieval patterns. We therefore used ERS to test the two hierarchically ordered predictions introduced above: first, that retrieval patterns would preferentially reinstate the absent reference point over the target; second, and more stringently, that trial-wise similarity to the reference-to-target path, rather than to either endpoint alone, would track judgment accuracy.

### Posterior parietal retrieval patterns were more similar to the reference point than to the target

To test the first prediction, we examined whether cue-evoked retrieval patterns preferentially expressed the absent reference point relative to both the remaining sequence positions and the cue-specified target. To do so, we correlated the cue-evoked pattern on each retrieval trial with the encoding pattern at each of the nine positions in the corresponding sequence (Fig. 4a). Across the six retrieval trial types, the group-mean profiles showed a common local maximum at the reference-point position in PCu and AG (Fig. 4b; trial-type- and hemisphere-specific profiles are shown in Supplementary Fig. 3). Reference-point ERS did not vary significantly by retrieval trial type in any ROI (one-way repeated-measures ANOVAs, all p ≥ 0.641, ηG² ≤ 0.009).

**Fig. 4.**
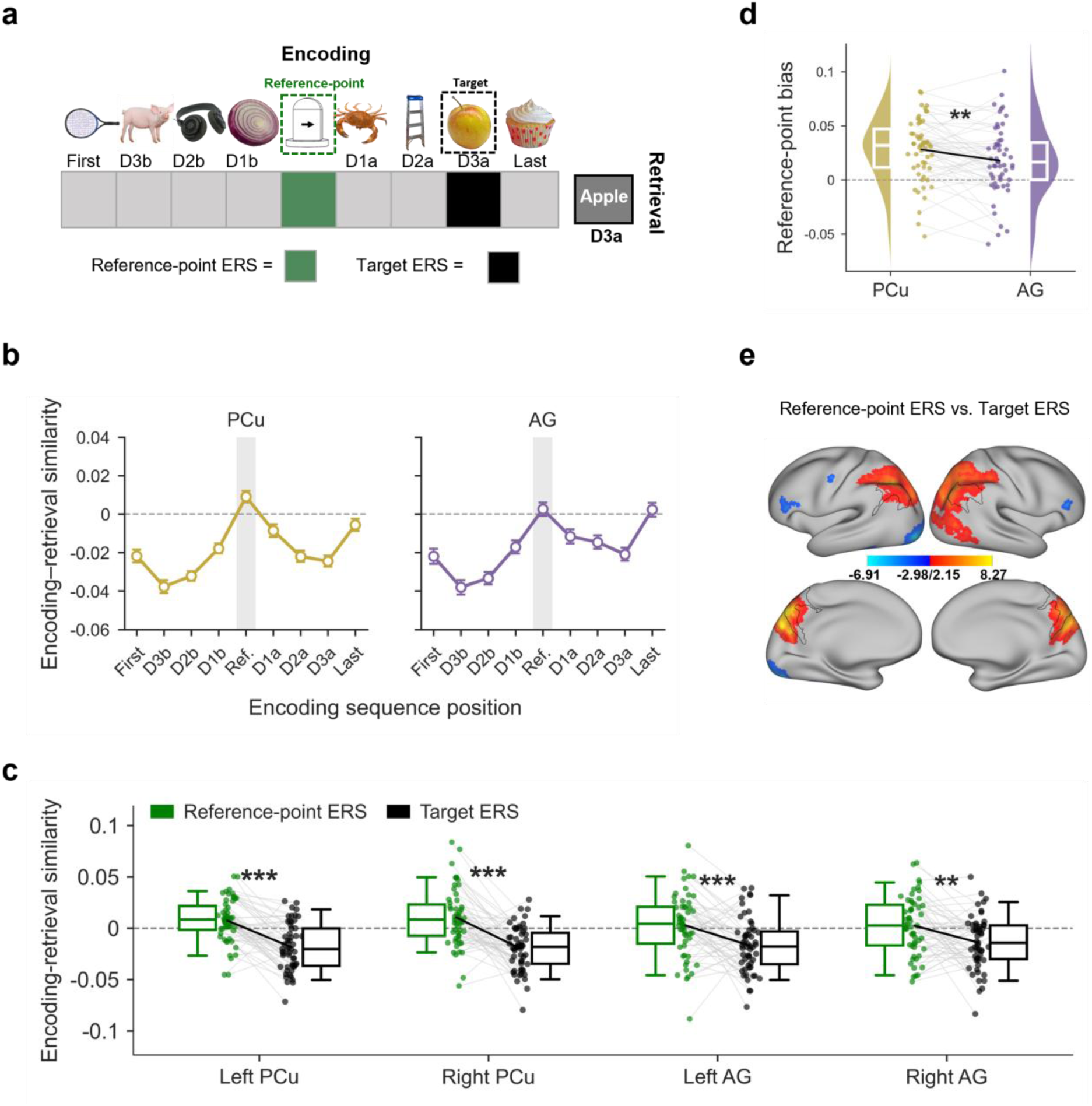
Encoding-position similarity and reference-point bias in posterior parietal retrieval patterns. **a,** Position-wise encoding–retrieval similarity (ERS) analysis. For each retrieval trial, the cue-evoked pattern was correlated with the encoding pattern at each of the nine positions in the same sequence. **b,** Mean position-wise ERS profiles in precuneus (PCu) and angular gyrus (AG), averaged across the six retrieval trial types and hemispheres. Points show across-participant means and error bars show s.e.m.; the shaded column marks the reference-point position. **c,** Reference-point ERS and target ERS in each ROI. Points show participant-level values, gray lines connect paired observations and boxplots show medians and interquartile ranges. **p_FDR_ < 0.01, ***p_FDR_ < 0.001, two-sided paired t tests. **d,** Reference-point bias (reference-point ERS minus target ERS), averaged across hemispheres, in PCu and AG. **p < 0.01, two-sided paired t test. **e,** Whole-brain searchlight contrast of reference-point ERS versus target ERS; warm and cool colors indicate positive and negative t values, respectively. Maps in e are displayed on inflated cortical surfaces; black contours mark the predefined PCu and AG ROIs. Significance was defined as p_FWE_ < 0.05, TFCE-corrected using non-parametric permutation testing.

We first compared reference-point ERS with the mean ERS for the seven same-sequence positions excluding the reference point and target. Reference-point ERS was higher than other-objects ERS in left PCu (t(54) = 9.07, p_FDR_ < 0.001, Hedges’ g = 1.21), right PCu (t(54) = 7.92, p_FDR_ < 0.001, Hedges’ g = 1.05), left AG (t(54) = 5.16, p_FDR_ < 0.001, Hedges’ g = 0.69), and right AG (t(54) = 5.48, p_FDR_ < 0.001, Hedges’ g = 0.73; Supplementary Fig. 4 and Supplementary Table 5). Thus, the reference-point maximum exceeded similarity to the remaining non-target sequence positions.

The theoretically critical comparison tested whether reference-point ERS also exceeded similarity to the cue-specified target. Reference-point ERS was higher than target ERS in left PCu (t(54) = 6.53, p_FDR_ < 0.001), right PCu (t(54) = 6.46, p_FDR_ < 0.001), left AG (t(54) = 3.64, p_FDR_ < 0.001), and right AG (t(54) = 3.35, p_FDR_ = 0.001; two-sided paired-samples t tests; Fig. 4c and Supplementary Table 6). This pattern shows a relative retrieval-pattern bias toward the absent reference point over the externally specified target.

We next tested whether the reference-point bias differed between PCu and AG. A 2 × 2 repeated-measures ANOVA on reference-point ERS minus target ERS, with brain region and hemisphere as within-participant factors, showed a main effect of brain region (F(1, 54) = 7.31, p = 0.009, ηG² = 0.021; Fig. 4d). The bias was greater in PCu than in AG (t(54) = 2.70, p = 0.009, Hedges’ g = 0.33). Neither the main effect of hemisphere nor the brain region × hemisphere interaction was significant (both p > 0.2). Component analyses showed higher reference-point ERS in PCu than AG (brain-region effect: F(1, 54) = 5.50, p = 0.023, p_FDR_ = 0.046, ηG² = 0.014; paired comparison: t(54) = 2.34, p = 0.023, Hedges’ g = 0.26), whereas no brain-region or hemisphere effects were detected for target ERS (all p > 0.2; Supplementary Table 7). The regional difference in the bias was therefore driven by its reference-point component.

A whole-brain searchlight analysis identified a bilateral posterior parietal cluster for the reference-point ERS > target ERS contrast, encompassing PCu, superior parietal cortex, and inferior parietal regions including AG (Fig. 4e and Supplementary Table 8). Additional clusters were located in right middle occipital and middle/inferior temporal cortices. The reverse contrast, target ERS > reference-point ERS, identified clusters in left middle/inferior occipital cortex, left fusiform/inferior temporal cortex, bilateral inferior frontal gyri, and left precentral/postcentral cortex (p_FWE_ < 0.05, TFCE-corrected).

Across the ROI analyses, retrieval patterns were more similar to the reference point than to both the remaining non-target sequence positions and the cue-specified target. The reference-point-minus-target difference was larger in PCu than in AG, and the same contrast identified an extended bilateral posterior parietal effect in the whole-brain analysis.

### Reference-to-target path ERS was associated with successful TOJ in bilateral PCu and AG

To test the second prediction, we examined whether trial-wise similarity to the encoded reference-to-target sequence segment was associated with successful TOJ. Reference-to-target path ERS was defined as the average similarity between the retrieval pattern and the encoding patterns of the milestone, target, and any intervening items along the current path (Fig. 5; see Methods for details).

**Fig. 5.**
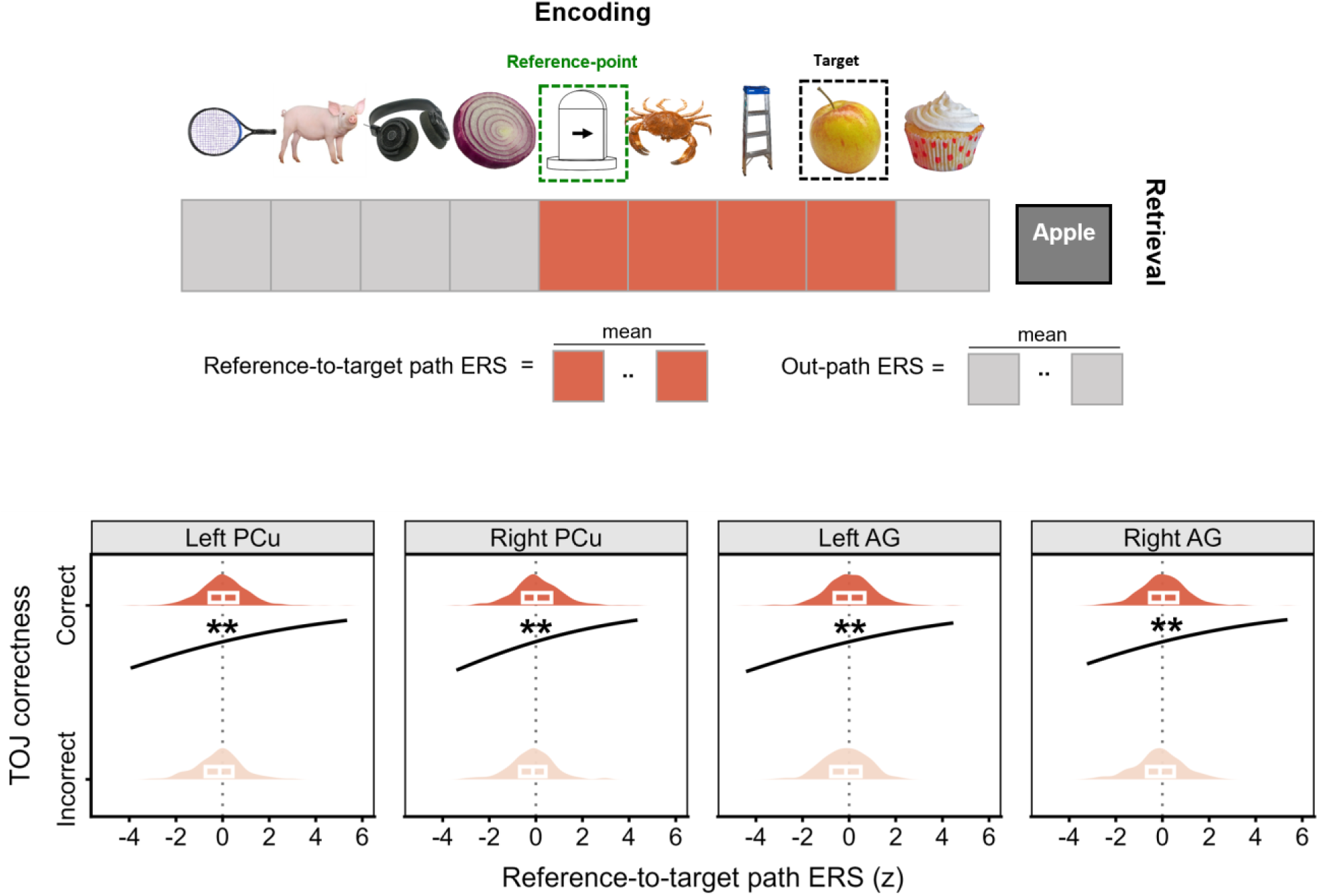
Reference-to-target path encoding–retrieval similarity and temporal-order judgment correctness. **Top,** Reference-to-target path ERS was the mean similarity between the retrieval pattern and encoding patterns at all positions from the reference point to the cue-specified target, including intervening items; out-path positions are shown in gray. **Bottom,** Mixed-effects logistic regression relating trial-wise path ERS to temporal-order judgment (TOJ) correctness in each region of interest (ROI). Black curves show population-level model-predicted probabilities of a correct TOJ, adjusted for trial type. Marginal density plots and boxplots show path-ERS distributions for correct and incorrect trials; boxplots show medians and interquartile ranges. **p_FDR_ < 0.01.

Reference-to-target path ERS was positively associated with trial-wise TOJ correctness in all four ROIs (Fig. 5; Supplementary Table 9): left PCu (z = 2.70, p_FDR_ = 0.007; odds ratio = 1.22, 95% CI = [1.06, 1.41]), right PCu (z = 3.31, p_FDR_ = 0.004; odds ratio = 1.28, 95% CI = [1.11, 1.48]), left AG (z = 2.69, p_FDR_ = 0.007; odds ratio = 1.22, 95% CI = [1.06, 1.41]), and right AG (z = 2.87, p_FDR_ = 0.007; odds ratio = 1.23, 95% CI = [1.07, 1.42]). The association remained significant in all four ROIs after controlling for retrieval activation (path ERS: z range = 2.82 to 3.42; all p_FDR_ ≤ 0.005; Supplementary Table 10).

To assess whether PCu and AG showed independent associations, we computed path ERS using merged left–right masks for PCu and AG and entered the two bilateral estimates simultaneously into a mixed-effects logistic regression model. After mutual adjustment, neither predictor showed an independent association with trial-wise TOJ correctness (PCu: z = 1.43, p = 0.152; AG: z = 1.57, p = 0.116; VIFs < 2).

As a complementary whole-brain searchlight analysis, we tested whether the ROI-level association between target-excluded path ERS and trial-wise TOJ correctness extended beyond the predefined ROIs. No searchlights survived whole-brain FDR correction (q < 0.05).

### Control analyses supported the specificity of the reference-to-target path ERS association

The second prediction further implied that the path ERS–correctness association should not be explained by similarity to either endpoint alone or to positions outside the task-relevant path. We therefore conducted control analyses addressing endpoint similarity, out-path similarity, and averaging-related properties of the ERS measure.

Because path ERS averaged similarity across the reference point, the target, and any intervening items, we first asked whether either endpoint measure alone could account for the path ERS–correctness association. Neither endpoint measure was significantly associated with trial-wise TOJ correctness in any ROI. Reference-point ERS estimates were positive in all four ROIs but were not significant (p ≥ 0.057; p_FDR_ ≥ 0.116; Supplementary Table 11), and target ERS likewise showed no significant association (p ≥ 0.102; p_FDR_ ≥ 0.153; Supplementary Table 12).

Second, because the reference point was repeated across sequences, we tested whether the path ERS association could be attributed to increased pattern reliability from repeated exposure, rather than to the reference point’s role within the task-relevant path. Under a repetition-only account, excluding the reference point should substantially reduce or eliminate the association. We therefore recomputed path ERS after excluding the reference point. Reference-excluded path ERS showed nominally significant positive associations in left PCu (p = 0.030), right PCu (p = 0.021), and right AG (p = 0.049), but not in left AG (p = 0.089). None of these associations survived FDR correction (p_FDR_ = 0.061–0.089; Supplementary Table 13). Because the reference point was an integral component of the task-relevant path, removing it was expected to weaken the association. This control therefore weakens, but does not eliminate, a repetition-only account, and the full reference-to-target path measure remained the measure most strongly associated with correctness.

Third, because the cue-specified target was perceptually present at retrieval, we tested whether the path ERS–correctness association depended on including it in the ERS measure. Target-excluded path ERS, defined as the average similarity between the retrieval pattern and the encoding patterns of the reference point and any intervening items while omitting the target, remained significantly associated with trial-wise TOJ correctness in all four ROIs: left PCu (z = 2.26, p_FDR_ = 0.024; odds ratio = 1.18, 95% CI = [1.02, 1.37]), right PCu (z = 3.20, p_FDR_ = 0.004; odds ratio = 1.27, 95% CI = [1.10, 1.47]), left AG (z = 2.41, p_FDR_ = 0.022; odds ratio = 1.19, 95% CI = [1.03, 1.38]), and right AG (z = 3.07, p_FDR_ = 0.004; odds ratio = 1.25, 95% CI = [1.08, 1.45]; Supplementary Table 14). This indicates that the path ERS–correctness association did not depend on perceptual reactivation of the cue-specified target.

Fourth, we tested whether the path ERS–correctness association reflected broader similarity to other sequence positions. Out-path ERS was defined as the average similarity between the retrieval pattern and encoding patterns of items outside the current reference-to-target path. When entered as the sole neural predictor in mixed-effects logistic regression models, out-path ERS was not significantly associated with trial-wise TOJ correctness in any ROI (all p_FDR_ ≥ 0.836; Supplementary Table 15). We then entered path ERS and out-path ERS simultaneously into the same models. Path ERS remained positively associated with trial-wise TOJ correctness after accounting for out-path ERS, whereas out-path ERS showed no independent association after accounting for path ERS (path ERS: z range = 2.70 to 3.25, all p_FDR_ ≤ 0.007; out-path ERS: all p_FDR_ ≥ 0.858; Supplementary Table 16). The same pattern was observed after additionally controlling for retrieval activation (path ERS: z range = 2.84 to 3.34, all p_FDR_ ≤ 0.005; out-path ERS: all p_FDR_ ≥ 0.322; Supplementary Table 17).

Fifth, we tested whether the path ERS effect could be explained by the greater reliability expected from averaging multiple encoding patterns. To match the number of encoding patterns contributing to the ERS measure, we generated length-matched pseudo-path ERS values. For each trial, pseudo-paths were created by sampling the same number of encoding positions as the actual reference-to-target path, but from positions outside the task-relevant path. This procedure was repeated 5,000 times. In each iteration, actual path ERS and length-matched pseudo-path ERS were entered simultaneously into a mixed-effects logistic regression model predicting trial-wise TOJ correctness. Across iterations, actual path ERS estimates were consistently positive in all four predefined ROIs. In contrast, the pseudo-path z-value distributions were centered near zero and their empirical 95% intervals overlapped zero (Supplementary Fig. 5). This analysis therefore provides a robustness characterization rather than an additional independent inferential test.

## Discussion

The central contribution of this study is content-specific representational evidence consistent with a reconstructive account of temporal-order retrieval. By separating the item that initiated retrieval from the temporal anchor needed to evaluate it, we could test two hierarchically ordered predictions: that retrieval patterns would preferentially express the absent, internally required reference point, and that successful judgments would be associated with similarity to the encoded information linking that anchor to the cue-specified target. Two findings addressed these predictions. First, retrieval patterns were more similar to the absent reference point than to either the cue-specified target or the remaining non-target positions, indicating a relative prioritization of the internally required anchor; this reference-point bias was stronger in the precuneus (PCu) than in the angular gyrus (AG). Second, trial-wise similarity to the encoded reference-to-target sequence segment was associated with correct judgments in both regions, even after accounting for retrieval activation. Together, these findings distinguish retrieval-related engagement from retrieval content: univariate activation provided an index of retrieval-related engagement in posterior parietal regions, whereas encoding–retrieval similarity characterized which encoding-related information was preferentially expressed within their retrieval patterns. Thus, the results are consistent with temporal-order retrieval being organized in relation to the relational information required for the judgment, rather than solely by the information specified by the external cue.

Reconstructive accounts hold that temporal judgments depend on active, repeated construction rather than on a fixed temporal code ^3,4^. Testing this prediction has been difficult, however, because in conventional temporal-order judgment tasks both compared items are typically presented at retrieval, so similarity to their encoding patterns can reflect cue processing as much as mnemonic recovery. Here, the target was specified by its name, whereas the milestone serving as the reference point was absent during retrieval, so any bias toward it could not reflect concurrent perceptual presentation. Retrieval patterns were indeed more closely aligned with the absent reference point than with either the cue-specified target or the remaining non-target positions—not a broad increase in similarity to all sequence items, but a bias specifically toward the absent milestone. This pattern was specific to posterior parietal cortex: the reverse contrast, favoring the target, was instead observed in occipital and fusiform regions associated with visual processing, consistent with perceptual reinstatement of the named target being confined to visual cortex rather than driving posterior parietal retrieval patterns. This regional dissociation argues against an account in which posterior parietal retrieval patterns specifically are governed by the cue’s perceptual properties.

The reference-point bias is also difficult to attribute to associative mechanisms that scale with distance between studied items, whether distance-graded spreading through a temporal-context representation ^11–13^ or simple contiguity-based association, strongest for adjacent items and weakening with lag ^30^. Both predict stronger reinstatement when the milestone is adjacent to the target (D1) than when intervening items separate the two (D2/D3). Contrary to this, reference-point ERS did not differ significantly across the six trial types spanning D1–D3. Nor is the bias readily explained by deliberate associative encoding, such as a narrative linking the milestone to other items: participants received no instruction to adopt such strategies, which typically require explicit prompting to be effective ^31^. This pattern is more consistent with selective orientation toward a task-defined anchor than with associative spreading or elaborative strategy, and aligns with the attention-to-memory (AtoM) model, in which dorsal parietal regions such as the precuneus support goal-directed, top-down orienting toward mnemonic information rather than being governed by the cue or the material’s associative structure ^25,26^.

Reference-point prioritization alone, however, cannot explain successful judgment: recovering the milestone identifies an anchor, but successful judgment additionally requires information that relates the target to that anchor. The path ERS analysis supplies a more direct and consequential link for reconstructive accounts. Reference-to-target path ERS captured similarity to the sequence segment spanning the reference point, the target, and any intervening items. Path ERS was positively associated with trial-wise correctness, whereas neither reference-point ERS nor target ERS showed a detectable association when examined alone. The bias identifies how retrieval content was oriented; path ERS identifies the relationally defined portion of the sequence whose expression tracked successful inference. Together they turn a general reconstructive proposal into a content-specific representational result: temporal judgments were accompanied by recovery of encoding-related information from a relationally defined sequence segment, consistent with accounts in which such judgments are derived from recovered episodic content rather than read from a stored timestamp ^3,4^, and extending evidence that information tied to intervening events contributes to temporal-order memory ^7^ to a reference-defined segment in human posterior parietal retrieval patterns. This association survived several specificity checks, including exclusion of the perceptually present target from the path measure, arguing against a purely perceptual account (Supplementary Tables 11–17). The representational basis of path ERS cannot be fully specified from the present data: because similarity was averaged across positions, the effect may reflect reactivation of individual path items, recovery of the temporal-contextual state linking them ^12,32^, or an integrated, holistic representation of the relevant sequence segment. This ambiguity does not, however, bear on the interpretation of the path ERS–correctness association as evidence for reconstruction: reconstructive accounts hold that temporal judgments depend on recovering useful information to infer position, not on any particular representational format, and each of these candidate mechanisms constitutes exactly this kind of recovered, task-relevant information.

These results provide representational content for PCu’s association with temporal reconstruction. Prior human neuroimaging showed that PCu activity varies with reconstruction demands, temporal relations among remembered events, and narrative context ^18,20,21^, but regional activation alone could not identify the mnemonic content being accessed. The present design closes that gap: PCu retrieval patterns were relatively aligned with an absent temporal reference point, and similarity to the reference-to-target segment covaried with judgment accuracy. The convergence with the univariate results reinforces this point—PCu activation showed the most robust independent association with performance, while PCu patterns also showed the stronger reference-point bias and a behaviorally relevant path-ERS association—indicating that the path-ERS association could not be reduced to retrieval-activation magnitude alone. These results also extend recent macaque recordings linking encoding–retrieval population similarity in medial posterior parietal cortex to successful temporal-order judgment ^22^: whereas that study identified a broad population-level link, the present design specifies which encoded content carries the behaviorally relevant correspondence in humans, suggesting cross-species convergence in the involvement of medial posterior parietal cortex in temporal-order retrieval.

The regional comparison was graded rather than categorical. Like PCu, but not strictly to the same degree, AG patterns were more similar to the reference point than to the target and remaining non-target positions, and AG path ERS was associated with trial-wise correctness—consistent with proposals that AG represents recovered episodic content and relational features of remembered events ^18,20,23,24^. Reference-point bias was weaker in AG, however; AG activation did not retain an independent association with correctness once PCu activation was included in the same model, and when PCu and AG path ERS were entered simultaneously, neither region retained an independent effect. This pattern does not support a strict functional dissociation between PCu and AG: both regions expressed task-relevant encoding–retrieval correspondence, and the loss of independent effects in the joint models may reflect shared variance between the two regions rather than identical contributions. The reference-point bias was nonetheless more pronounced in PCu.

The AtoM framework distinguishes exactly this kind of dorsal, top-down orienting— consistent with the stronger reference-point bias observed in PCu—from a more ventral, bottom-up registration of retrieved content, associated with regions such as AG ^25,26^. These findings motivate a schematic model of reference-based TOJ (Fig. 6). In this model, PCu may play a stronger role in orienting mnemonic processing according to retrieval goals, whereas AG may contribute to representing or maintaining recovered temporal-relational information in interaction with dorsal parietal goal-directed processes. These complementary contributions may support temporal reconstruction by allowing the encoded relation between anchor and target to be inferred relative to the internal reference point. This schematic model is necessarily confined to posterior parietal cortex, however, leaving open how the proposed reconstruction relates to hippocampal contributions to temporal-order memory. Hippocampal activity patterns encode sequential and event-structure information during encoding and support their reinstatement at retrieval ^33–36^, and dynamically reorganize around event boundaries ^9^; posterior parietal cortex is, in turn, thought to interact with the hippocampus within a broader posterior-medial memory system supporting contextual and relational retrieval ^16,37^. The present analyses focused on the a priori posterior parietal regions motivated by prior work on temporal-order retrieval; testing whether the reference-point bias and path ERS reported here in PCu and AG depend on, parallel, or are downstream of hippocampal reinstatement of the encoded sequence is a natural and directly tractable extension— for example, by defining a hippocampal ROI within the same analysis pipeline and testing whether hippocampal and posterior parietal reference-to-target similarity covary on a trial-by-trial basis.

**Fig. 6.**
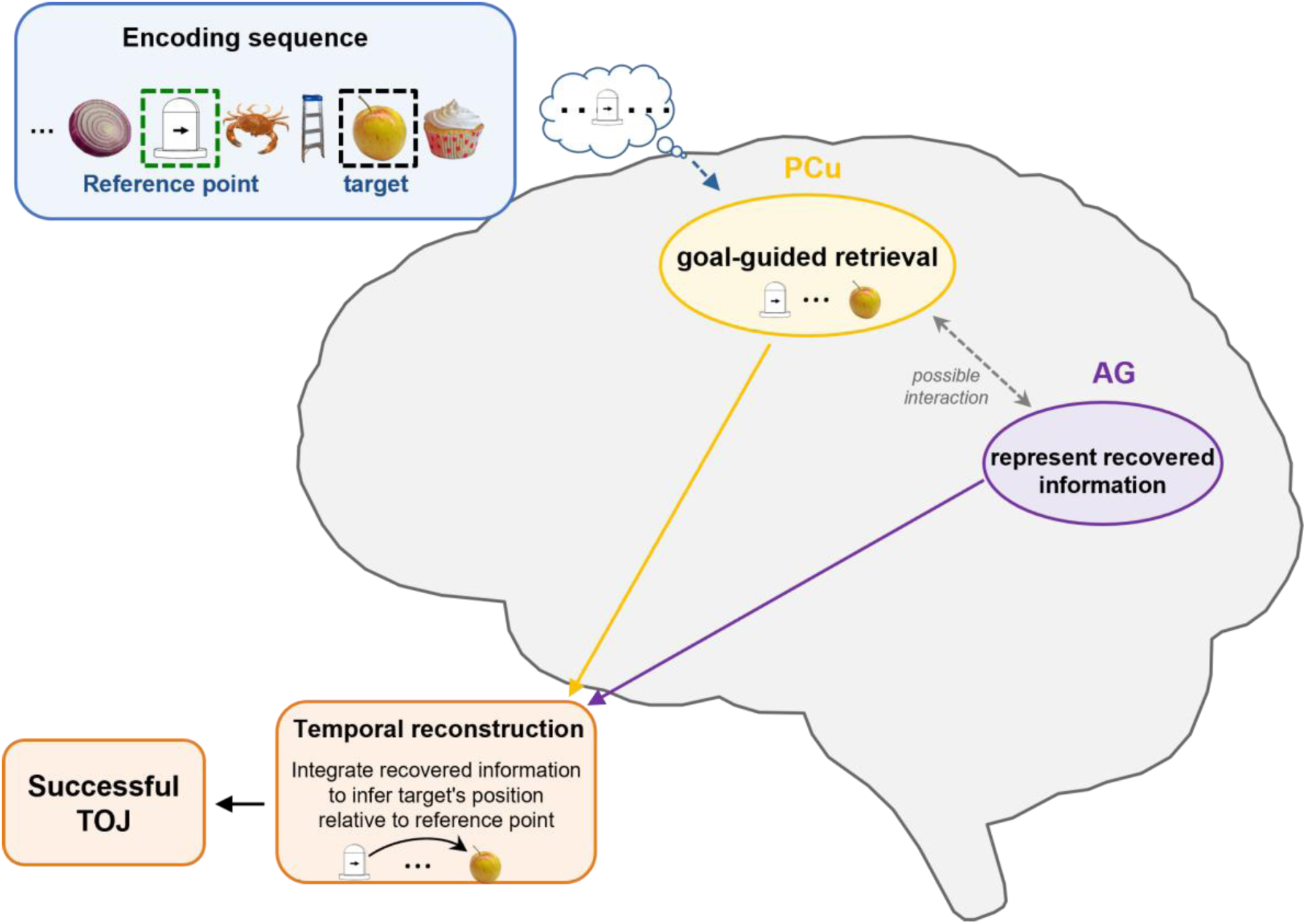
Proposed posterior parietal contributions to reference-based temporal-order judgment. The cue identifies the target, whereas the absent reference point and task-relevant reference-to-target information are proposed to be internally accessed to support the judgment. Precuneus (PCu) retrieval patterns showed stronger reference-point bias and the most robust retrieval-activation association with correctness; angular gyrus (AG) showed weaker reference-point bias but behaviorally relevant reference-to-target path ERS. Solid arrows summarize hypothesized contributions to temporal reconstruction, and the dashed bidirectional arrow denotes possible PCu–AG interaction. The schematic is a functional summary and does not imply temporal order, causality or directed information flow.

Several limitations should be noted. First, ERS measures correspondence between encoding and retrieval patterns, not the conscious recovery or causal use of specific memories, so it cannot establish whether PCu or AG—or their proposed differentiated roles in Fig. 6—are necessary for the judgment or engaged in a particular temporal order; perturbational and temporally resolved methods could address this directly. A further limitation concerns the milestone itself: as the task-defined reference point, the only repeated item, and a fixed serial position, its contribution to the reference-point bias and path-ERS association cannot be uniquely attributed to task relevance alone. Existing control analyses only partly constrain this ambiguity (Supplementary Tables 11, 13), and fully resolving it would require a design in which the same stimulus serves as an anchor for some judgments but not others, which we identify as a direction for future work. In addition, testing each encoded sequence only once—by design, to avoid retrieval-practice effects and carryover across retrieval episodes—limited the number of trial-level observations available per participant; future designs that preserve this single-shot logic while sampling more sequences per participant would sharpen individual-level brain–behavior estimates. Finally, the highly controlled sequences and fixed reference point isolate reference-defined temporal structure but limit generalization to natural memories, which involve multiple candidate anchors and unevenly salient episodes.

By dissociating the cue-specified target from the reference point required for judgment, this study turns a general reconstructive account into a content-specific representational test, and does so with a design simple enough to extend to other domains where retrieval must be organized around a goal rather than a cue. Reference-point bias shows that posterior parietal retrieval patterns were oriented toward an internally required anchor; path ERS links successful judgment to encoding-related information spanning the anchor-to-target relation. This evidence does not identify a replay mechanism or confine reconstruction to posterior parietal cortex, but it advances the human literature beyond activation-based inference by showing which encoded information is expressed during temporal-order retrieval and how it relates to behavior. In reference-based temporal-order judgment, these results suggest that determining when an event occurred relative to an internal anchor depends not only on recovering the event itself, but on configuring retrieval around the relational structure needed to place it in time. In this sense, the human precuneus does not simply reinstate the past; its retrieval states are selectively organized around the temporal structure required for the current mnemonic decision.

## Methods

### Participants

Fifty-seven healthy university students participated in the experiment. Two participants were excluded from all analyses because severe motion artifacts rendered their fMRI data unusable across the entire scan. The final sample therefore included 55 participants (15 women; mean age = 19.4 ± 1.5 years, range = 18–24 years). All participants had normal or corrected-to-normal vision. All experimental procedures were approved by the Ethics Committee of Kochi University of Technology, and all participants provided written informed consent before the experiment.

### Experimental stimuli

The stimulus set consisted of one milestone image, created with Microsoft PowerPoint, and 192 color images of nameable everyday objects. The milestone image served as the fixed temporal reference point across sequences and was not used as a retrieval cue; retrieval was initiated by the name of a target object. The object images were selected from a standardized stimulus bank ^38^ and covered a broad range of object categories to reduce reliance on strong pre-existing semantic associations. All images were resized to 500 × 500 pixels. The 192 objects were pseudo-randomly assigned to four runs and further divided into six blocks within each run, with eight objects per block. Both stimulus order within blocks and block order were randomized for each participant.

### Experimental procedure

Fig. 1 illustrates the overall experimental design. The experiment was conducted using PsychoPy v2023.2.3 ^39^, with a total scanning time of approximately 45 min. Participants received task instructions and completed a brief practice run outside the scanner. After the practice session, they entered the scanner and completed four experimental runs. Each run contained six blocks, and each block consisted of three phases: encoding, interference, and retrieval.

The encoding phase began with a fixation cross shown for 4.5 s. One milestone image and eight unique object images were then presented sequentially against a gray background (RGB: [160, 160, 160]), with each image subtending a visual angle of 2°. The milestone always appeared in the middle of the sequence, at the fifth position, whereas the positions of the eight object images were randomized for each participant. Each image was displayed for 1.5 s, with a fixed interstimulus interval (ISI) of 4.5 s. Participants were instructed to silently name each object.

A 10-s interference phase was included after encoding to reduce conscious rehearsal of the encoded stimuli. The phase consisted of a self-paced odd–even judgment task on numbers below 100. Participants pressed the left button for odd numbers and the right button for even numbers, with numbers presented in a randomized order and separated by a 0.5-s intertrial interval. They were instructed to prioritize response accuracy.

The retrieval phase began with a 4.5-s fixation cross, followed by the name of one encoded object for 1.5 s. When the object-name cue appeared, participants retrieved the object’s temporal position relative to the milestone during encoding. After a 4.5-s delay, a response timeline was shown for up to 4 s, with the milestone fixed at the center. Participants reported the remembered position as quickly and accurately as possible by moving a slider with the left and right buttons. The slider started at the center of the timeline, and each button press moved it by one discrete position to the left or right. The timeline remained on screen until participants confirmed their response with the up button or until the 4-s response window elapsed. Because objects had been presented as images during encoding, object names, rather than object images, were used as retrieval cues to reduce perceptual overlap between encoding and retrieval ^40,41^.

Each retrieval trial was assigned to one of six trial types according to the cued object’s temporal distance (TD) from the milestone and whether it appeared before or after the milestone during encoding. TD was operationalized as the positional distance between the cued object and the milestone in the encoding sequence, such that TD = 1 indicated no intervening objects, TD = 2 indicated one intervening object, and TD = 3 indicated two intervening objects. Specifically, D1b, D2b, and D3b referred to objects presented before the milestone at TDs of 1, 2, and 3, respectively; D1a, D2a, and D3a referred to objects presented after the milestone at TDs of 1, 2, and 3, respectively. Objects presented first (D4b) or last (D4a) during encoding were excluded from retrieval to avoid potential primacy and recency effects ^42,43^. Each block contained one retrieval trial, and the six blocks within each run corresponded to the six trial types. This design ensured that each encoded sequence was tested only once, thereby reducing potential retrieval-practice effects and minimizing carryover from prior retrieval episodes ^44,45^.

After the scanning session, participants completed an exploratory post-task questionnaire indicating which temporal position they perceived as easiest to remember. Because participants could select more than one option, the questionnaire was treated as an exploratory measure of subjective memorability rather than a forced-choice ranking. The five response options were “Earlier,” “D1b,” “Same,” “Later,” and “D1a.” “Earlier” and “Later” indicated that target objects presented earlier or later in the sequence were perceived as easier to remember, respectively. “D1b” and “D1a” indicated that the objects immediately before or after the milestone were perceived as easier to remember, respectively. “Same” indicated no perceived difference across temporal positions.

### Run-level exclusions and retained trials

Among the 55 included participants, 13 runs across 9 participants were excluded from run-level analyses: 3 for excessive head motion (mean framewise displacement [FD] > 0.35 mm or > 50% of volumes classified as motion outliers), 6 for sleepiness, and 4 for visual obstruction caused by glasses fogging. The latter 10 exclusions were based on contemporaneous scanner-session records. Each participant with run-level exclusions retained at least two valid runs. After these exclusions, 46 of the 55 included participants retained the full set of 24 retrieval trials, corresponding to 4 trials for each of the 6 trial types. The remaining 9 participants retained 12 or 18 trials, corresponding to 2 or 3 trials per trial type.

### Behavioral statistical analysis

Behavioral analyses were conducted on the same 55 participants included in the fMRI analyses. Trial-wise TOJ correctness was averaged within each participant for each combination of relative position (before vs. after the milestone) and temporal distance (D1, D2, and D3), yielding condition-level accuracy estimates. Accuracy was analyzed using a 2 × 3 repeated-measures ANOVA with relative position and temporal distance as within-participant factors. Greenhouse–Geisser-corrected p values were reported when sphericity was violated. Significant interactions were followed by Bonferroni-corrected simple-effects comparisons within before- and after-milestone trials. Effect sizes for ANOVAs were reported as generalized eta squared, and paired comparisons were reported with Hedges’ g ^46,47^.

Because multiple selections were allowed in the exploratory post-task questionnaire, questionnaire responses were analyzed using participant-level permutation tests. For each permutation, the number of options selected by each participant was preserved, but selected options were randomly reassigned across the five response categories. An overall chi-square-type statistic tested whether the observed response distribution deviated from the permutation null distribution. Follow-up one-sided permutation tests assessed whether each response option was selected more often than expected under the same null model, with Bonferroni correction applied across the five options.

### MRI acquisition and preprocessing

Neuroimaging data were collected using a Siemens Magnetom Prisma 3T MRI scanner equipped with a 64-channel head coil. Functional images were acquired using a multiband echo-planar imaging (EPI) sequence (repetition time [TR] = 743 ms; echo time [TE] = 36.5 ms; flip angle [FA] = 48°; field of view [FOV] = 192 mm; matrix = 96 × 96; in-plane resolution = 2 × 2 mm²; slice thickness = 2 mm; 72 slices; multiband acceleration factor = 8). A high-resolution whole-brain structural image was acquired using a 3D T1-weighted MPRAGE sequence (TR = 1900 ms; TE = 2.52 ms; FA = 8°; FOV = 250 mm; matrix = 256 × 256; in-plane resolution = 1 × 1 mm²; slice thickness = 1 mm; 176 slices).

All raw MRI data were converted to Brain Imaging Data Structure (BIDS, version 1.0.1) format and validated using the BIDS Validator. Data were preprocessed using fMRIPrep 25.1.1, which is built on Nipype 1.10.0.

Structural preprocessing began with intensity non-uniformity correction of the T1-weighted image using N4BiasFieldCorrection implemented in ANTs ^48,49^, and the corrected image was used to generate the T1w reference. The T1w reference was skull-stripped using an ANTs-based brain-extraction workflow, and brain tissue segmentation into cerebrospinal fluid (CSF), white matter (WM), and gray matter (GM) was performed using FSL FAST ^50^. Cortical surface reconstruction was performed with FreeSurfer version 7.3.2 ^51^, after which the T1w reference was nonlinearly normalized to the MNI152NLin6Asym ^52^ template using ANTs.

To correct susceptibility-induced distortions in the functional data, fieldmap-less susceptibility distortion correction implemented in SDCFlows was applied. In this procedure, a deformation field was estimated by co-registering the EPI reference image to the participant’s T1w reference ^53,54^ using antsRegistration (ANTs 2.6.0), with deformation constrained to the phase-encoding direction.

Functional preprocessing was performed separately for each run. The first six volumes were discarded to allow for signal stabilization. A reference volume was generated for motion correction, and head-motion parameters were estimated concurrently. Functional images were co-registered to the T1w anatomical reference using boundary-based registration ^55^ and subsequently resampled to the specified MNI standard space. Several nuisance regressors were computed, including framewise displacement (FD), DVARS ^56^, and global signals from CSF, WM, and the whole brain. Physiological noise components were further estimated using both temporal CompCor (tCompCor) and anatomical CompCor (aCompCor) ^57^. Specifically, tCompCor components were derived from the top 2% most variable voxels within the brain mask, whereas aCompCor components were extracted from WM and CSF masks, retaining enough components to explain 50% of the variance within those masks. Motion-outlier volumes were identified by fMRIPrep based on FD > 0.5 mm or standardized DVARS > 1.5.

Additional preprocessing was performed before statistical analyses. For whole-brain univariate analyses, standard-space functional images were spatially smoothed using a 6-mm full-width-at-half-maximum (FWHM) Gaussian kernel implemented in FSL SUSAN ^58^ and temporally high-pass filtered with a 100-s cutoff. For ROI-based univariate and ERS analyses, unsmoothed native-space functional data were high-pass filtered using the same 100-s cutoff. For searchlight ERS analyses, unsmoothed standard-space functional data were high-pass filtered using the same 100-s cutoff.

### ROI definition

Four a priori ROIs (Fig. 3a) were defined in each participant’s native T1 space using FreeSurfer cortical parcellation tools (https://surfer.nmr.mgh.harvard.edu/) and the Destrieux atlas ^59^. The ROIs included bilateral precuneus (PCu) and bilateral angular gyrus (AG), corresponding to the Destrieux labels G_precuneus and G_pariet_inf-Angular, respectively. For ROI-based univariate activation analyses and encoding–retrieval similarity analyses, the FreeSurfer-derived ROI masks were resampled to the functional image grid and used to extract unsmoothed functional data. For visualization, the corresponding Destrieux atlas labels were displayed on a standard cortical surface using Connectome Workbench (version 2.1.0) ^60^.

### Univariate analysis

First-level models were specified in FEAT and estimated using FILM prewhitening in FSL version 6.0.6 ^61,62^. Encoding object events and retrieval cue events were modeled as separate conditions. For the main contrast, encoding object events corresponding to objects that were later tested during retrieval were included in the encoding condition of interest, and the corresponding retrieval cue events were included in the retrieval condition of interest. Thus, the retrieval > encoding contrast compared retrieval cue-evoked activity with encoding object-evoked activity for the same set of tested items. Untested encoding object events, milestone events, interference-task periods, and button-press events during retrieval were modeled as regressors of no interest. Encoding object events and retrieval cue events of interest were modeled at stimulus onset with a duration of 1.5 s. All task regressors were convolved with the double-gamma hemodynamic response function, and temporal derivatives were included to improve model sensitivity. Nuisance regressors included six head-motion parameters and six aCompCor components. In addition, censor regressors were added for motion-outlier volumes identified by fMRIPrep. For each censor regressor, the corresponding outlier time point was coded as 1 and all other time points as 0, allowing the GLM to account for transient motion-related artifacts. Each run was modeled separately in the first-level analysis. The same first-level GLM was used for both analyses, providing β estimates for the ROI-based percent-signal-change (PSC) analysis and retrieval > encoding contrast images for the whole-brain analysis at the run level.

For the ROI-based PSC analysis, β estimates for the encoding and retrieval regressors of interest were extracted from the first-level GLM estimates within each predefined ROI and averaged across voxels within each ROI. PSC was calculated within each run as (β / mean signal intensity) × ppheight × 100%, where ppheight represents the peak height of the hemodynamic response relative to baseline ^63^. Run-level PSC values were averaged across runs within each participant before group-level statistical testing. Paired-samples t tests were then used to compare PSC between retrieval and encoding phases within each ROI. Statistical significance was controlled for multiple comparisons across the four predefined ROIs using FDR correction ^64^, and effect sizes were reported as Hedges’ g. To compare the magnitude of retrieval-minus-encoding activation between PCu and AG, retrieval-minus-encoding PSC differences were also submitted to a 2 × 2 repeated-measures ANOVA with brain region (PCu vs. AG) and hemisphere (left vs. right) as within-participant factors.

To characterize whole-brain retrieval-related engagement, first-level contrast images were computed for the retrieval > encoding contrast in each run. These run-level contrast images were combined within each participant using a fixed-effects model, yielding one participant-level contrast image per participant. Group-level whole-brain inference was then performed using non-parametric permutation testing implemented in FSL randomise ^65^. Specifically, participant-level contrast images were merged into a 4D image and tested against zero using a one-sample sign-flipping test with 10,000 permutations. Multiple comparisons were controlled using threshold-free cluster enhancement (TFCE) ^66^, and statistical significance was defined as p_FWE_ < 0.05.

### Trial-specific activity patterns

Preprocessed functional data were entered into a nuisance-only voxel-wise general linear model (GLM) implemented in FSL FEAT. Six head-motion parameters, six aCompCor components, and censor regressors for motion-outlier volumes were included as nuisance predictors. Only the residuals of this GLM, defined as the portion of the signal not explained by these nuisance regressors, were retained for subsequent ERS analyses ^34,36^. Voxel-wise residual time series were extracted from each predefined ROI and normalized by z-scoring within each voxel ^27^. Because task timings were defined in multiples of the TR, each encoding event, including object and milestone presentations, and each retrieval event, defined as target-name cue presentation, was aligned with an fMRI volume onset. The onset-aligned volume was indexed as volume 0. Trial-specific activity patterns were estimated by averaging volumes 5, 6, and 7 after this onset-indexed volume. Given the TR of 743 ms, the onsets of these volumes occurred approximately 3.7, 4.5, and 5.2 s after the indexed volume, and their acquisition windows spanned approximately 3.7–5.9 s. This window therefore covered approximately the 4–6 s post-onset hemodynamic response period and was selected to capture the expected peak response for the current event ^67^.

For encoding events, this window occurred after the preceding stimulus and before the onset of the subsequent stimulus. Because successive stimulus onsets were separated by 6 s (1.5-s stimulus presentation plus a 4.5-s ISI), these volumes were acquired approximately 10–12 s after the onset of the preceding stimulus, by which time the associated BOLD response was expected to have largely returned toward baseline. This timing reduced, but may not have fully eliminated, contamination from adjacent encoding events. For retrieval events, this window fell within the cue/delay period (1.5-s cue presentation followed by a 4.5-s delay) and ended before the onset of the timeline response screen, on which the milestone was displayed. Therefore, retrieval patterns estimated from this window could not reflect direct visual input from the milestone displayed on the response screen.

### Encoding–retrieval similarity analysis

Encoding–retrieval similarity (ERS) analysis ^27,28,68^ was conducted to quantify similarity between encoding-phase representations and retrieval patterns. In each ROI, the retrieval activity pattern for each trial was correlated with the encoding activity patterns from the same sequence. The resulting correlation coefficients were Fisher z-transformed before statistical analysis.

For each retrieval trial, the retrieval activity pattern was correlated separately with the encoding activity pattern at each of the nine positions in the corresponding sequence. Reference-point ERS was the similarity to the milestone at the fifth position, and target ERS was the similarity to the cue-specified target position. Other-objects ERS was the mean similarity to the seven same-sequence positions excluding both the reference point and target.

For each ROI, a one-way repeated-measures ANOVA tested whether reference-point ERS differed among the six target conditions. Planned two-sided paired-samples t tests compared reference-point ERS with other-objects ERS and with target ERS; FDR correction was applied across the four ROIs separately for each comparison. To test regional differences in the theoretically critical reference-point-versus-target effect, reference-point-minus-target ERS scores were entered into a 2 × 2 repeated-measures ANOVA with brain region (PCu vs. AG) and hemisphere (left vs. right) as within-participant factors. The bias was then averaged across hemispheres within each region and compared between PCu and AG using a paired-samples t test. To characterize the source of the regional bias difference, reference-point ERS and target ERS were analyzed separately using the same 2 × 2 ANOVA structure; the two planned brain-region p values were FDR-corrected. Hedges’ g was reported for paired comparisons and generalized eta squared for ANOVAs.

To characterize the spatial distribution of the reference-point-versus-target effect beyond the predefined ROIs, we conducted a whole-brain searchlight ERS analysis ^69^ in standard space using unsmoothed data. For each participant and run, trial-specific encoding and retrieval activity patterns were extracted within a 5 × 5 × 5 voxel searchlight centered on each gray-matter voxel. Within each searchlight, reference-point ERS and target ERS were computed using the same procedure as in the ROI analysis and averaged across valid runs for each participant. Participant-specific reference-point-minus-target ERS maps were entered into FSL randomise and tested against zero using a one-sample sign-flipping test with 10,000 permutations and threshold-free cluster enhancement. Statistical significance was defined as p_FWE_ < 0.05, TFCE-corrected. The reverse contrast was tested using sign-reversed difference maps.

### Mixed-effects logistic regression of brain–behavior relationships

Mixed-effects logistic regression models, implemented as binomial generalized linear mixed-effects models with a logit link, were used to test associations between trial-wise neural measures and trial-wise TOJ correctness. Trial-wise TOJ correctness was modeled as a binary dependent variable (correct = 1, incorrect = 0). Models were fitted separately for each predefined ROI. Each model included the neural measure of interest and trial type as fixed effects, with trial type (D3b, D2b, D1b, D1a, D2a, and D3a) modeled as an additional categorical factor to control for condition-level performance differences. A participant-specific random intercept accounted for repeated observations within participants.

For the activation analyses, encoding activation and retrieval activation were defined separately. Encoding activation was defined as the mean value across ROI voxels in the trial-specific encoding activity pattern described above for the cued target item. Retrieval activation was defined as the mean value across ROI voxels in the trial-specific retrieval activity pattern described above, which was extracted before the onset of the response screen. Separate activation models were specified as: Correctness ∼ encodingActivation + TrialType + (1 | Subject) and Correctness ∼ retrievalActivation + TrialType + (1 | Subject). As a complementary simultaneous-region analysis, bilateral PCu retrieval activation and bilateral AG retrieval activation were entered simultaneously into the same model to assess whether each region showed an independent association with trial-wise TOJ correctness after adjusting for the other region: Correctness ∼ PCu_retrievalActivation + AG_retrievalActivation + TrialType + (1 | Subject).

For the ERS analyses, reference-point ERS, target ERS, path ERS, and out-path ERS were defined separately. Path ERS was defined as the average similarity between the retrieval activity pattern and the encoding patterns of all positions included in the reference-to-target path, including the milestone, the target, and any intervening items. This measure was adapted from the encoding– retrieval neural global pattern similarity (ER-nGPS) approach ^70^, which quantifies the similarity between a retrieval activity pattern and the set of encoding activity patterns corresponding to studied items. Here, we adapted this logic to temporal-order retrieval by using path ERS to capture similarity to the reference-to-target sequence segment needed for the current judgment. Out-path ERS was defined as the average similarity between the retrieval activity pattern and the encoding patterns of same-sequence items that were not part of the current reference-to-target path.

To test whether path ERS was associated with trial-wise TOJ correctness, the main model was: Correctness ∼ pathERS + TrialType + (1 | Subject). To test whether the path ERS effect was independent of overall retrieval activation, retrieval activation was added as an additional fixed-effect predictor: Correctness ∼ pathERS + retrievalActivation + TrialType + (1 | Subject). As a complementary simultaneous-region analysis, bilateral PCu path ERS and bilateral AG path ERS were entered simultaneously into the same model to assess whether each region showed an independent association with trial-wise TOJ correctness after adjusting for the other region: Correctness ∼ PCu_pathERS + AG_pathERS + TrialType + (1 | Subject).

For the simultaneous-region PCu/AG models, bilateral PCu and bilateral AG masks were created by merging the corresponding left- and right-hemisphere ROI masks. Retrieval activation and path ERS predictors were recomputed within each bilateral mask rather than averaged across unilateral ROI estimates. Variance inflation factors (VIFs) were inspected to assess multicollinearity between PCu and AG predictors. VIFs were modest for the activation model (PCu = 1.94; AG = 1.94) and the path ERS model (PCu = 1.63; AG = 1.61), indicating acceptable collinearity. These simultaneous-region models assessed independent associations after mutual adjustment and were not formal tests of differences in coefficient magnitude.

Several control analyses were conducted separately for each of the four predefined ROIs to assess alternative explanations for the path ERS effect. First, to examine whether the effect could be explained by endpoint representations, separate models tested reference-point ERS and target ERS as predictors of trial-wise TOJ correctness: Correctness ∼ referenceERS + TrialType + (1 | Subject) and Correctness ∼ targetERS + TrialType + (1 | Subject). Second, because the same milestone image was repeated across sequences whereas target objects were unique, we recomputed path ERS after excluding the reference point from the reference-to-target path. Reference-excluded path ERS was then tested as a predictor of trial-wise TOJ correctness: Correctness ∼ referenceExcludedPathERS + TrialType + (1 | Subject). Third, because the target was specified by name at retrieval, we recomputed path ERS after excluding the target item. Target-excluded path ERS was defined as the average similarity between the retrieval pattern and the encoding patterns of the reference point and any intervening items, and was tested as a predictor of trial-wise TOJ correctness: Correctness ∼ targetExcludedPathERS + TrialType + (1 | Subject). Fourth, to assess whether out-path ERS itself was associated with trial-wise TOJ correctness, out-path ERS was entered as the sole neural predictor: Correctness ∼ outPathERS + TrialType + (1 | Subject). We then tested whether the path ERS association was independent of out-path similarity by entering path ERS and out-path ERS into the same model: Correctness ∼ pathERS + outPathERS + TrialType + (1 | Subject). A further control model additionally included retrieval activation: Correctness ∼ pathERS + outPathERS + retrievalActivation + TrialType + (1 | Subject). Finally, to test whether the path ERS effect could be explained by the greater reliability expected when multiple encoding patterns are averaged, we performed a length-matched pseudo-path control analysis. For each trial in each iteration, length-matched pseudo-path ERS was generated by randomly sampling the same number of encoding positions as included in the actual reference-to-target path from out-path positions within the same sequence. This procedure was repeated 5,000 times. In each iteration, actual path ERS and pseudo-path ERS were entered simultaneously into a mixed-effects logistic regression model with trial-wise TOJ correctness as the binary outcome: Correctness ∼ pathERS + pseudoPathERS + TrialType + (1 | Subject). The distributions of coefficient estimates and nominal p values across iterations were summarized, including the proportions of iterations in which each coefficient was positive and nominally significant (p < 0.05).

Continuous neural predictors were first mean-centered within participants and then z-scored across all trials before model fitting. This procedure improved model convergence and yielded standardized coefficients for the brain–behavior models. Standardized neural measures were then entered as continuous fixed-effect predictors in the mixed-effects logistic regression models. All models were fitted in R version 4.5.2 using the lme4 package ^71^. Model parameters were estimated by maximum likelihood using the BOBYQA optimizer. Statistical significance of fixed effects was evaluated using Wald z tests. Regression coefficients (β), odds ratios, 95% confidence intervals, and p values were reported for brain–behavior models. For separate ROI-level brain–behavior models, p values were FDR-corrected across the four predefined ROIs for each planned model and neural predictor.

Whole-brain searchlight analyses were conducted as complementary, exploratory analyses of the spatial distribution of brain–behavior relationships. For each trial, neural measures were extracted from cubic searchlights centered on gray-matter voxels. At each searchlight location, trial-wise TOJ correctness was modeled using the same mixed-effects logistic regression framework as in the ROI analyses. For each searchlight sphere, the neural measure was mean-centered within participants, z-scored across all retained trials, and entered as the predictor of interest. Trial type was included as an additional fixed effect, and a random intercept for participant accounted for repeated observations within participants. For the path ERS searchlight analysis, the cue-specified target was excluded from the path average, leaving the reference point and any intervening positions. Retrieval-activation and target-excluded path-ERS searchlight maps were corrected for multiple comparisons across searchlights using FDR correction at q < 0.05. For retrieval-activation results, only clusters containing at least 20 voxels were reported.

## Data availability

Processed data will be made available in a public repository before publication.

## Code availability

Analysis code will be made available in a public repository before publication.

## Author contributions

C.H. and K.N. conceived the project. C.H., J.F. and K.N. collected data. C.H. and R.W. analyzed data. C.H., J.F., R.W., I.H., K.J. and K.N. wrote the manuscript.

## Acknowledgments

We would like to thank Ms. Maoko Yamanaka for her administrative assistance. This study was supported by KAKENHI from the Japan Society for the Promotion of Science (23H00413 to K.N. and I.H.), and by AMED under grant number JP26wm0625205 to K.N. and I.H.

---

**Supplementary Fig. 1.**
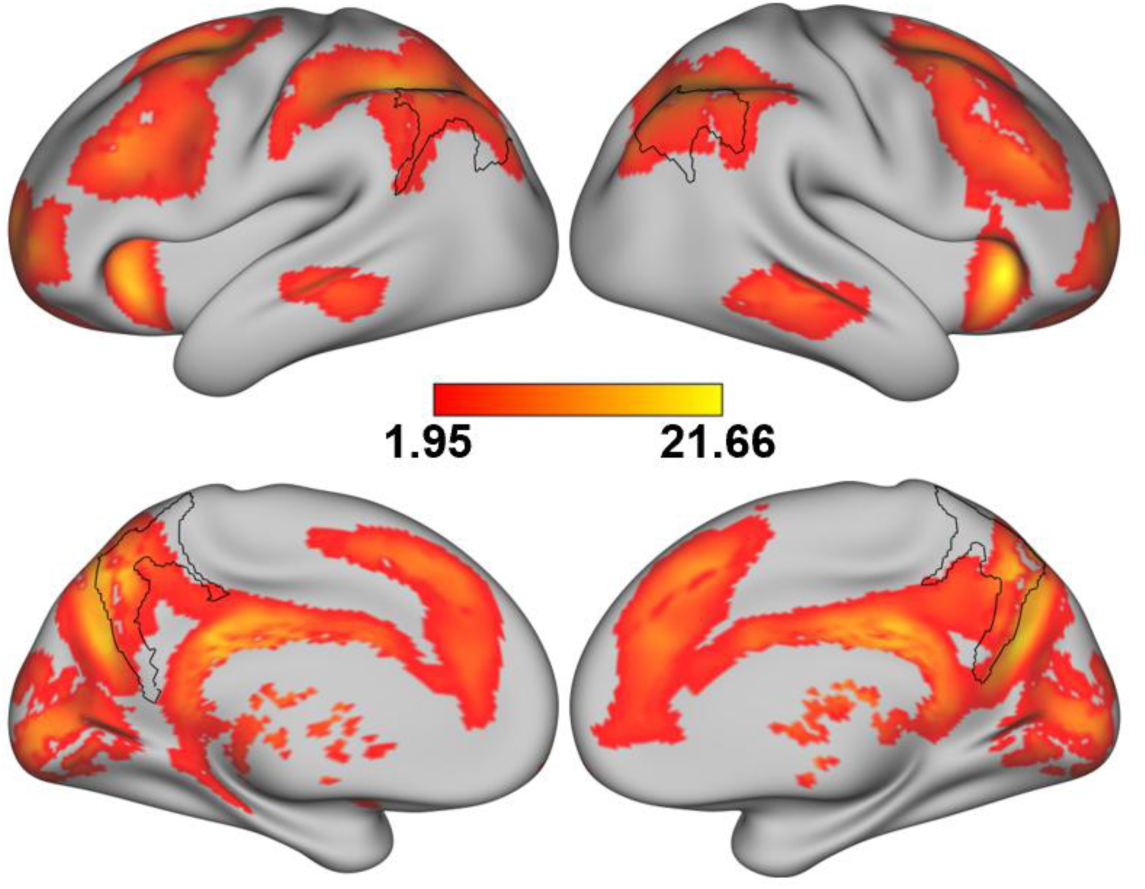
Whole-brain retrieval > encoding activation. Statistical maps show the retrieval > encoding contrast on inflated cortical surfaces. Black contours mark predefined bilateral precuneus (PCu) and angular gyrus (AG) regions of interest (ROIs). Significance was defined as p_FWE_ < 0.05, TFCE-corrected using non-parametric permutation testing; only clusters of at least 20 voxels are displayed. The color bar indicates t values.

**Supplementary Fig. 2.**
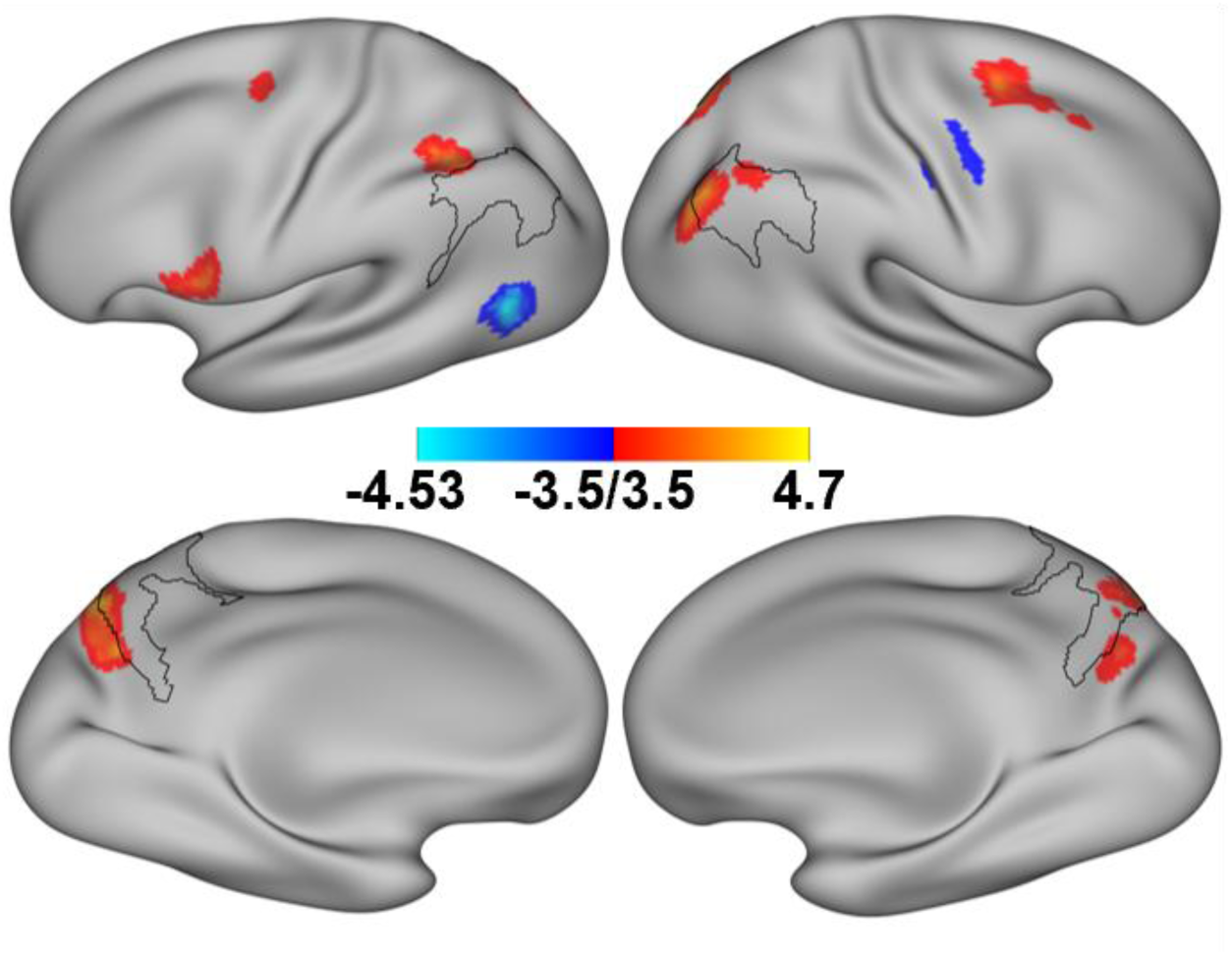
Whole-brain searchlight associations between retrieval activation and trial-wise temporal-order judgment correctness. Fixed effect of cue-evoked retrieval activation from searchlight-wise mixed-effects logistic regression models. At each gray-matter searchlight, temporal-order judgment correctness was modeled with retrieval activation as the predictor of interest, trial type as a categorical fixed effect and participant as a random intercept. Results were FDR-corrected across searchlights at q < 0.05. Warm and cool colors indicate positive and negative Wald z values, respectively. Only clusters containing at least 20 voxels are displayed. Black contours mark predefined bilateral precuneus (PCu) and angular gyrus (AG) regions of interest.

**Supplementary Fig. 3.**
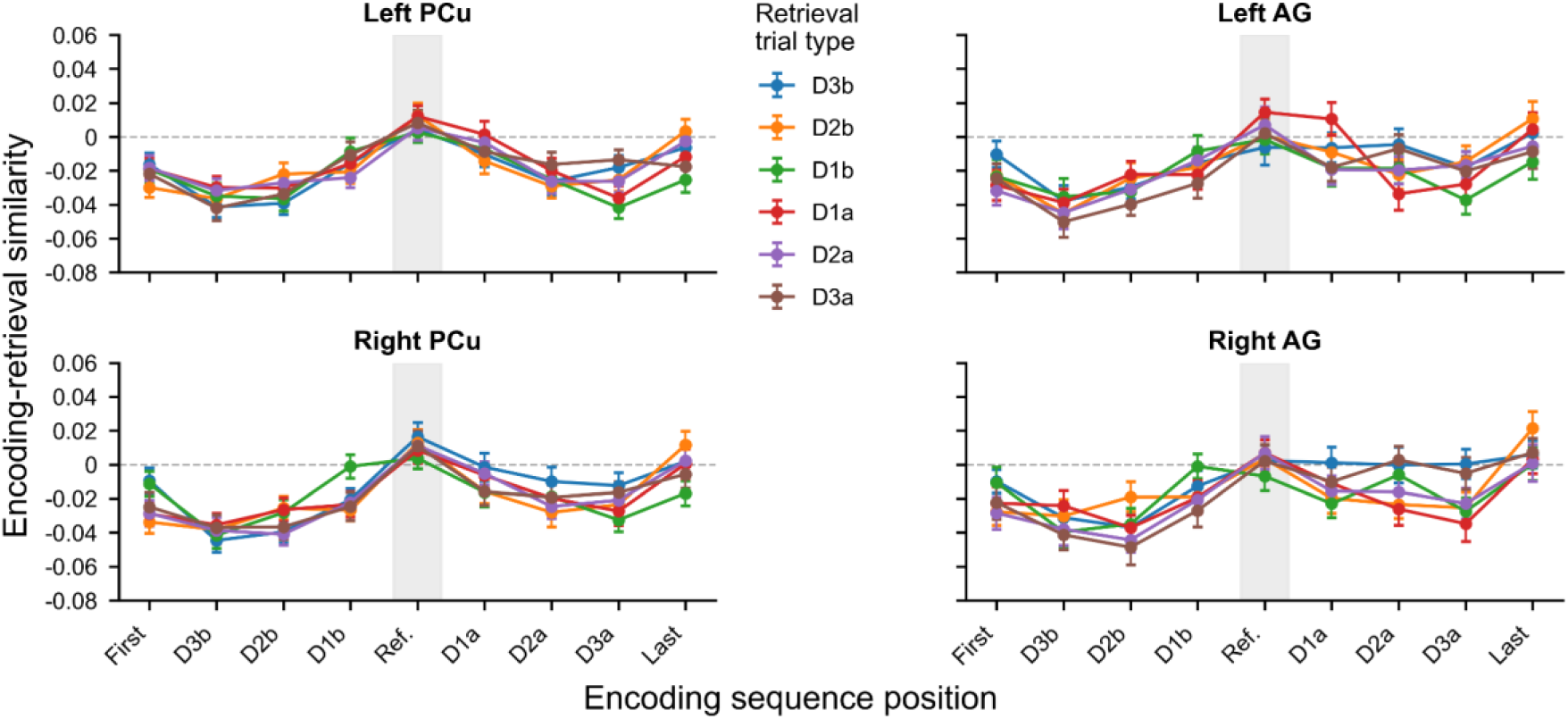
Position-wise encoding–retrieval similarity profiles by retrieval trial type and hemisphere. Mean position-wise ERS profiles are shown separately for the six retrieval trial types in left and right precuneus (PCu) and angular gyrus (AG). Points indicate across-participant means and error bars indicate s.e.m. The shaded column marks the reference-point position.

**Supplementary Fig. 4.**
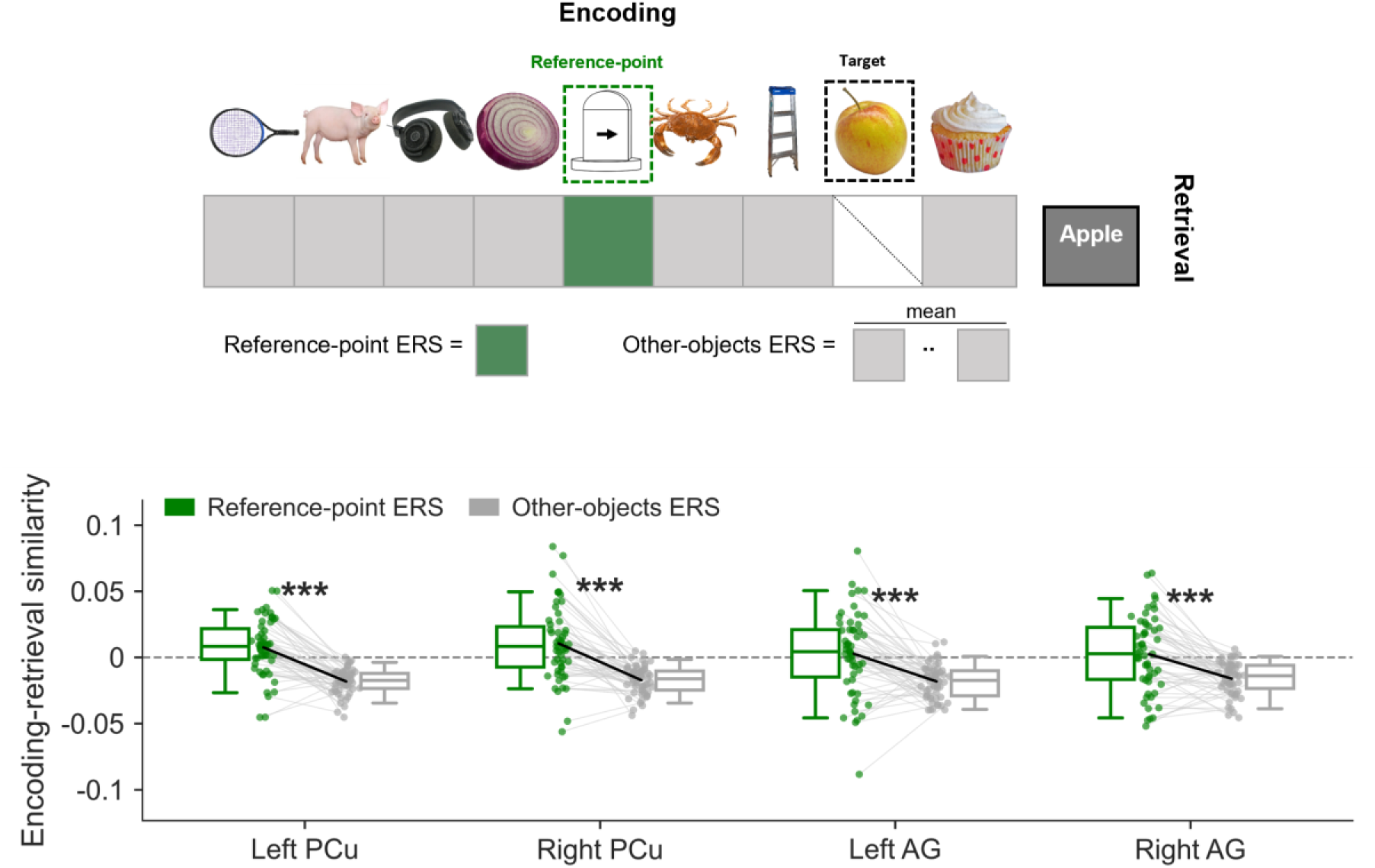
Reference-point and other-objects encoding–retrieval similarity. Top, Other-objects ERS was the mean similarity between each retrieval pattern and encoding patterns at the seven same-sequence positions excluding the reference point and cue-specified target. **Bottom,** Reference-point ERS and other-objects ERS in bilateral precuneus (PCu) and angular gyrus (AG). Points show participant-level values, gray lines connect paired observations and boxplots show medians and interquartile ranges. Asterisks denote two-sided paired t tests with FDR correction across the four regions of interest (***p_FDR_ < 0.001).

**Supplementary Fig. 5.**
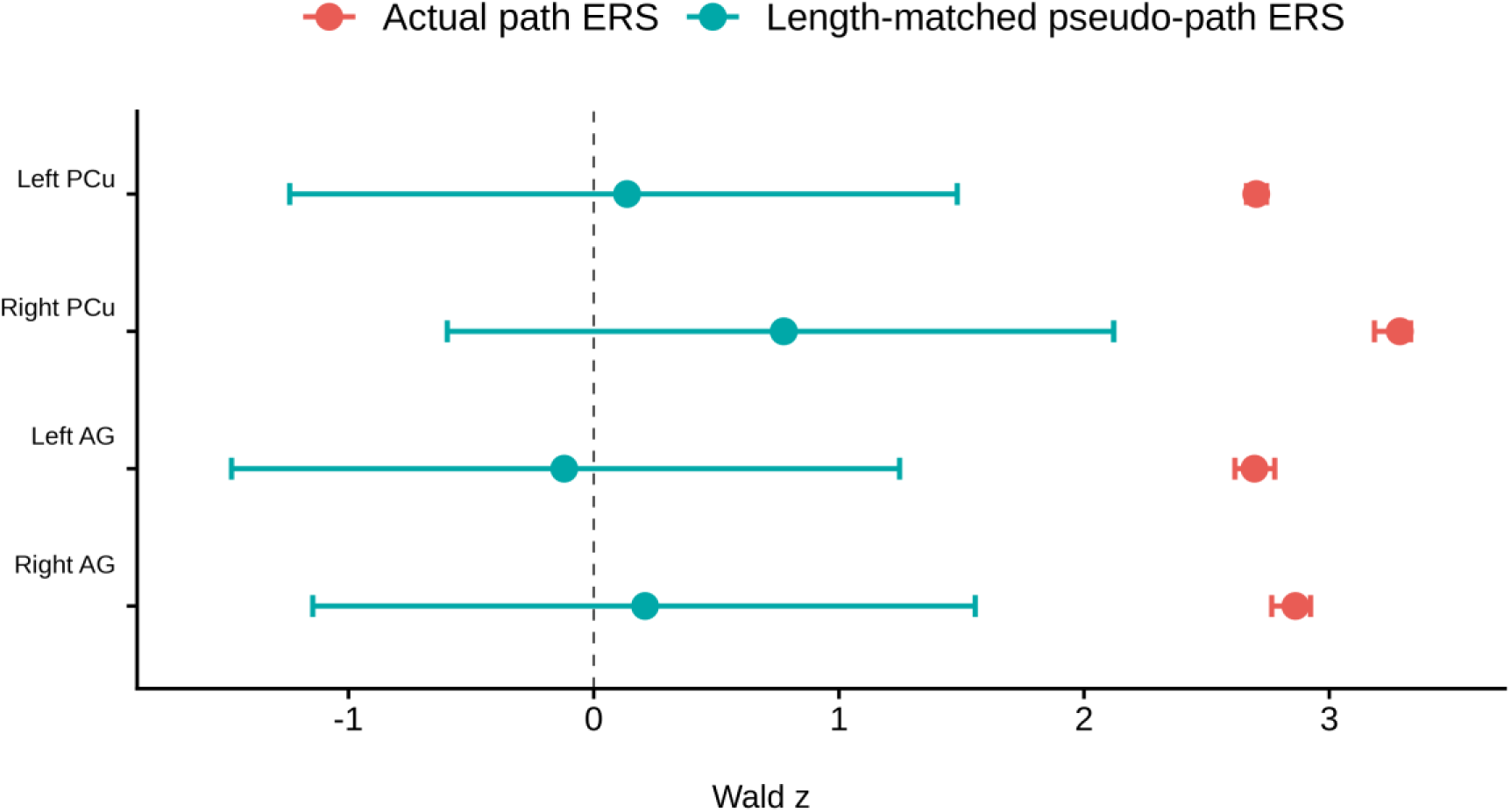
Length-matched pseudo-path control for reference-to-target path encoding– retrieval similarity. Dots show median Wald z values across 5,000 mixed-effects logistic regression models; horizontal lines show empirical 95% intervals (2.5th–97.5th percentiles). Each model included actual reference-to-target path ERS and one length-matched pseudo-path ERS measure as simultaneous predictors of trial-wise temporal-order judgment correctness. Pseudo-paths sampled the same number of encoding positions as the actual path from out-path positions in the same sequence. The dashed vertical line marks z = 0. Results are shown for the four predefined regions of interest using data from *n* = 55 participants.

**Whole-brain activation results for retrieval > encoding**

**Supplementary Table 1.** AAL-region summaries of the whole-brain retrieval > encoding contrast.

| Brain region | Hemi. | MNI coordinates |  |  | Peak<br>t | Voxel<br>count |
| --- | --- | --- | --- | --- | --- | --- |
|  |  | x | y | z |  |  |
| Parietal |  |  |  |  |  |  |
| Precuneus | L | -10 | -60 | 14 | 15.81 | 1539 |
| Precuneus | R | 30 | -50 | 8 | 17.62 | 1256 |
| Parietal_Inf | L | -34 | -74 | 42 | 12.43 | 1494 |
| Parietal_Inf | R | 58 | -56 | 40 | 17.48 | 958 |
| Angular | L | -38 | -56 | 32 | 12.60 | 609 |
| Angular | R | 44 | -68 | 30 | 16.75 | 890 |
| Parietal_Sup | L | -14 | -72 | 40 | 13.15 | 702 |
| Parietal_Sup | R | 32 | -68 | 50 | 12.55 | 979 |
| SupraMarginal | L | -58 | -28 | 22 | 4.23 | 134 |
| SupraMarginal | R | 60 | -46 | 24 | 12.23 | 319 |
| Frontal |  |  |  |  |  |  |
| Frontal_Inf_Tri | L | -52 | 28 | -2 | 20.89 | 912 |
| Frontal_Inf_Tri | R | 50 | 46 | 0 | 11.44 | 340 |
| Frontal_Inf_Orb_2 | L | -42 | 16 | -10 | 16.37 | 177 |
| Frontal_Inf_Orb_2 | R | 54 | 24 | -12 | 12.12 | 51 |
| Frontal_Sup_2 | L | -22 | 66 | 4 | 10.14 | 959 |
| Frontal_Sup_2 | R | 16 | 66 | 18 | 14.32 | 1795 |
| Frontal_Mid_2 | L | -40 | 50 | 4 | 11.31 | 2032 |
| Frontal_Mid_2 | R | 48 | 44 | 20 | 13.23 | 1553 |
| Frontal_Inf_Oper | L | -52 | 14 | 6 | 7.34 | 151 |
| Frontal_Inf_Oper | R | 44 | 16 | 10 | 11.26 | 512 |
| Frontal_Sup_Medial | L | -6 | 66 | 12 | 10.97 | 901 |
| Frontal_Sup_Medial | R | 12 | 54 | 2 | 10.20 | 185 |
| OFCant | L | -22 | 34 | -18 | 7.92 | 121 |
| OFCant | R | 34 | 42 | -20 | 5.67 | 58 |
| OFCmed | R | 20 | 24 | -26 | 3.97 | 37 |
| Medial temporal |  |  |  |  |  |  |
| Hippocampus | R | 34 | -6 | -26 | 6.24 | 39 |
| Occipital |  |  |  |  |  |  |
| Cuneus | L | -4 | -86 | 22 | 20.13 | 733 |
| Cuneus | R | 16 | -86 | 18 | 16.30 | 514 |
| Occipital_Sup | L | -12 | -102 | 12 | 17.55 | 297 |
| Occipital_Sup | R | 30 | -74 | 18 | 11.33 | 450 |
| Occipital_Mid | L | -14 | -96 | 0 | 10.60 | 371 |
| Occipital_Mid | R | 36 | -96 | 4 | 11.46 | 338 |
| Calcarine | L | 0 | -82 | -4 | 10.30 | 1299 |
| Calcarine | R | 18 | -94 | -4 | 10.95 | 643 |
| Lingual | L | -18 | -56 | -12 | 9.91 | 577 |
| Lingual | R | 14 | -88 | -12 | 10.59 | 540 |
| Fusiform | L | -20 | 0 | -46 | 6.14 | 25 |
| Occipital_Inf | L | -46 | -76 | -16 | 5.49 | 30 |
| **Temporal** |  |  |  |  |  |  |
| Temporal_Mid | L | -54 | -18 | -24 | 8.21 | 837 |
| Temporal_Mid | R | 56 | -10 | -24 | 7.12 | 376 |
| Temporal_Inf | L | -40 | 12 | -40 | 6.14 | 271 |
| Temporal_Inf | R | 36 | 8 | -46 | 4.92 | 100 |
| Temporal_Sup | R | 46 | -6 | -16 | 3.75 | 29 |
| **Insula** |  |  |  |  |  |  |
| Insula | L | -36 | -8 | -10 | 21.66 | 344 |
| Insula | R | 26 | 18 | -16 | 15.56 | 294 |
| **Cingulate and medial cortex** |  |  |  |  |  |  |
| Cingulate_Post | L | -12 | -40 | 12 | 11.74 | 188 |
| Cingulate_Post | R | 4 | -44 | 12 | 15.75 | 63 |
| Cingulate_Mid | L | -6 | -32 | 32 | 10.29 | 533 |
| Cingulate_Mid | R | 2 | -32 | 30 | 12.89 | 450 |
| ACC_sup | L | -8 | 28 | 22 | 9.59 | 359 |
| ACC_sup | R | 14 | 34 | 20 | 7.99 | 251 |
| ACC_pre | L | -4 | 40 | 12 | 8.30 | 423 |
| ACC_pre | R | 2 | 42 | 4 | 6.72 | 182 |
| **Sensorimotor** |  |  |  |  |  |  |
| Precentral | L | -46 | -2 | 32 | 10.83 | 548 |
| Precentral | R | 62 | 4 | 32 | 12.29 | 564 |
| Supp_Motor_Area | L | -4 | 16 | 44 | 12.02 | 426 |
| Supp_Motor_Area | R | 6 | 16 | 46 | 10.01 | 325 |
| Postcentral | L | -58 | -20 | 14 | 5.00 | 55 |
| Postcentral | R | 56 | -2 | 20 | 8.59 | 487 |
| **Subcortical** |  |  |  |  |  |  |
| Caudate | L | -6 | 12 | 12 | 8.75 | 511 |
| Caudate | R | 12 | 10 | 8 | 10.21 | 404 |
| Thal_PuM | L | -22 | -32 | 6 | 9.41 | 78 |
| Thal_PuM | R | 16 | -32 | 4 | 8.49 | 41 |
| Putamen | L | -30 | -8 | -8 | 6.05 | 129 |
| Putamen | R | 32 | 8 | -8 | 8.31 | 85 |
| N_Acc | R | 6 | 10 | -10 | 5.91 | 38 |
| **Cerebellum** |  |  |  |  |  |  |
| Cerebellum_7b | L | -44 | -64 | -54 | 14.26 | 159 |
| Cerebellum_7b | R | 48 | -52 | -56 | 14.38 | 140 |
| Cerebellum_Crus1 | L | -38 | -72 | -34 | 13.30 | 877 |
| Cerebellum_Crus1 | R | 36 | -46 | -36 | 13.87 | 836 |
| Cerebellum_8 | L | -16 | -56 | -54 | 8.08 | 458 |
| Cerebellum_8 | R | 34 | -58 | -60 | 13.87 | 238 |
| Cerebellum_6 | L | -28 | -42 | -30 | 12.32 | 674 |
| Cerebellum_6 | R | 42 | -34 | -32 | 13.77 | 526 |
| Cerebellum_Crus2 | L | -44 | -68 | -50 | 13.26 | 347 |
| Cerebellum_Crus2 | R | 50 | -62 | -46 | 11.52 | 484 |
| Vermis_7 |  | 6 | -78 | -28 | 12.53 | 79 |
| Cerebellum_10 | L | -14 | -36 | -44 | 12.37 | 65 |
| Cerebellum_10 | R | 24 | -36 | -44 | 8.87 | 55 |
| Vermis_9 |  | 2 | -58 | -38 | 11.97 | 76 |
| Vermis_10 |  | 4 | -42 | -40 | 10.86 | 22 |
| Cerebellum_9 | L | -2 | -54 | -52 | 10.24 | 226 |
| Cerebellum_9 | R | 12 | -42 | -54 | 8.49 | 184 |
| Vermis_4_5 |  | 6 | -54 | -24 | 8.95 | 147 |
| Vermis_3 |  | -2 | -42 | -20 | 8.90 | 65 |
| Vermis_8 |  | 0 | -64 | -44 | 8.70 | 25 |
| Cerebellum_3 | L | -12 | -40 | -26 | 6.88 | 26 |
| Cerebellum_4_5 | L | -24 | -44 | -30 | 6.86 | 262 |
| Cerebellum_4_5 | R | 30 | -34 | -36 | 5.14 | 23 |
| Vermis_6 |  | 4 | -58 | -24 | 6.79 | 162 |

*Note.* Only AAL3 regions with at least 20 suprathreshold voxels are reported, grouped by broad anatomical division and ordered by absolute peak t. AAL3 masks were applied to the TFCE-corrected map; peak coordinates identify the maximum absolute t in each region, and voxel count is the number of suprathreshold voxels in that region. Hemisphere is reported separately. AAL, Automated Anatomical Labeling; Hemi., hemisphere; MNI, Montreal Neurological Institute; TFCE, threshold-free cluster enhancement.

**Supplementary Table 2.** Retrieval activation predicting trial-wise temporal-order judgment correctness.

| ROI | Predictor | $\beta$ | z | Odds ratio | 95% CI | p | p<sub>FDR</sub> |
| --- | --- | --- | --- | --- | --- | --- | --- |
| Left PCu | retrieval activation | 0.22 | 3.00 | 1.24 | [1.08, 1.43] | 0.003 | **0.005\*\*** |
| Right PCu | retrieval activation | 0.30 | 4.10 | 1.35 | [1.17, 1.55] | < 0.001 | **< 0.001\*\*\*** |
| Left AG | retrieval activation | 0.18 | 2.51 | 1.20 | [1.04, 1.38] | 0.012 | **0.016\*** |
| Right AG | retrieval activation | 0.13 | 1.83 | 1.14 | [0.99, 1.31] | 0.068 | **0.068~** |

**Supplementary Table 3.** Encoding activation predicting trial-wise temporal-order judgment correctness.

| ROI | Predictor | $\beta$ | z | Odds ratio | 95% CI | p | p<sub>FDR</sub> |
| --- | --- | --- | --- | --- | --- | --- | --- |
| Left PCu | encoding activation | -0.03 | -0.37 | 0.97 | [0.85, 1.12] | 0.713 | 0.713 |
| Right PCu | encoding activation | 0.06 | 0.80 | 1.06 | [0.92, 1.22] | 0.424 | 0.566 |
| Left AG | encoding activation | 0.13 | 1.89 | 1.14 | [0.99, 1.32] | 0.059 | 0.237 |
| Right AG | encoding activation | 0.08 | 1.14 | 1.08 | [0.94, 1.25] | 0.254 | 0.507 |

**Supplementary Table 4.** Searchlight clusters in which retrieval activation was associated with trial-wise temporal-order judgment correctness.

| Cluster No. | Brain region | Hemi. | Peak MNI |  |  | Peak z | K-E (voxels) | AAL overlap |
| --- | --- | --- | --- | --- | --- | --- | --- | --- |
|  |  |  | x | y | z |  |  |  |
| Positive association with TOJ correctness |  |  |  |  |  |  |  |  |
| C1 | Precuneus / Superior parietal lobule | Bilateral | -10 | -72 | 44 | 4.72 | 409 | 45.97% Precuneus (L); 28.12% Precuneus (R); 17.60% Superior parietal lobule (R) |
| C2 | Angular gyrus / Middle occipital gyrus | R | 38 | -76 | 42 | 4.42 | 154 | 55.19% Angular gyrus (R); 33.77% Middle occipital gyrus (R); 7.79% Superior occipital gyrus (R) |
| C4 | Inferior parietal lobule | L | -38 | -50 | 40 | 4.24 | 53 | 98.11% Inferior parietal lobule (L) |
| C11 | Cerebellum Crus I | L | -42 | -42 | -34 | 4.23 | 23 | 100.00% Cerebellum Crus I (L) |
| C5 | Superior frontal gyrus / Middle frontal gyrus | R | 28 | 0 | 56 | 4.21 | 39 | 92.31% Superior frontal gyrus (R); 7.69% Middle frontal gyrus (R) |
| C6 | Inferior frontal gyrus, opercular part / Precentral gyrus | L | -54 | 8 | 14 | 4.11 | 36 | 77.78% Inferior frontal gyrus, opercular part (L); 22.22% Precentral gyrus (L) |
| C3 | Angular gyrus / Inferior parietal lobule | R | 44 | -58 | 50 | 4.10 | 54 | 75.93% Angular gyrus (R); 22.22% Inferior parietal lobule (R) |
| C7 | Precentral gyrus / Middle frontal gyrus | L | -22 | -10 | 46 | 4.10 | 34 | 55.88% no_label; 20.59% Precentral gyrus (L); 11.76% Middle frontal gyrus (L); 11.76% Superior frontal gyrus (L) |
| C8 | Precuneus | R | 16 | -58 | 30 | 3.99 | 32 | 96.88% Precuneus (R) |
| C10 | Middle frontal gyrus | R | 36 | 26 | 54 | 3.87 | 25 | 92.00% Middle frontal gyrus (R) |
| C9 | Middle frontal gyrus / Superior frontal gyrus | R | 34 | 14 | 62 | 3.87 | 25 | 52.00% Middle frontal gyrus (R); 36.00% Superior frontal gyrus (R); 12.00% no_label |
| Negative association with TOJ correctness |  |  |  |  |  |  |  |  |
| C1 | Middle occipital gyrus / Middle temporal gyrus | L | -44 | -68 | 6 | -4.54 | 75 | 54.67% Middle occipital gyrus (L); 45.33% Middle temporal gyrus (L) |
| C3 | Caudate | L | -8 | 8 | 6 | -4.50 | 30 | 100.00% Caudate (L) |
| C2 | Precentral gyrus / Middle frontal gyrus | R | 54 | -8 | 52 | -4.08 | 34 | 70.59% Precentral gyrus (R); 29.41% Middle frontal gyrus (R) |

*Note.* Each row represents one contiguous suprathreshold cluster from a searchlight mixed-effects logistic regression model of trial-wise temporal-order judgment correctness. Trial type was included as a categorical fixed effect and participant as a random intercept. Peak z values are signed Wald statistics for retrieval activation. Clusters are grouped by association direction and were FDR-corrected across searchlights at $q < 0.05$ ; only clusters of at least 20 voxels are reported. Cluster labels were assigned with AtlasReader using the AAL atlas and may include multiple AAL regions; AAL overlap includes all reported entries, including no\_label. AAL, Automated Anatomical Labeling; Hemi., hemisphere; MNI, Montreal Neurological Institute; K-E, cluster extent.

**Supplementary Table 5.** Reference-point ERS compared with other-objects ERS.

| ROI | Contrast | t | df | p<sub>FDR</sub> | Hedges' g |
| --- | --- | --- | --- | --- | --- |
| Left PCu | Reference-point ERS vs. other-objects ERS | 9.07 | 54 | <0.001 | 1.21 |
| Right PCu | Reference-point ERS vs. other-objects ERS | 7.92 | 54 | <0.001 | 1.05 |
| Left AG | Reference-point ERS vs. other-objects ERS | 5.16 | 54 | <0.001 | 0.69 |
| Right AG | Reference-point ERS vs. other-objects ERS | 5.48 | 54 | <0.001 | 0.73 |

*Note.* Two-sided paired-samples t tests compared reference-point ERS with mean similarity to the seven same-sequence positions excluding the reference point and cue-specified target. p values were FDR-corrected across the four regions of interest (all p<sub>FDR</sub> < 0.001). Hedges' g is the paired standardized mean difference with small-sample correction. ERS, encoding–retrieval similarity.

**Supplementary Table 6.**
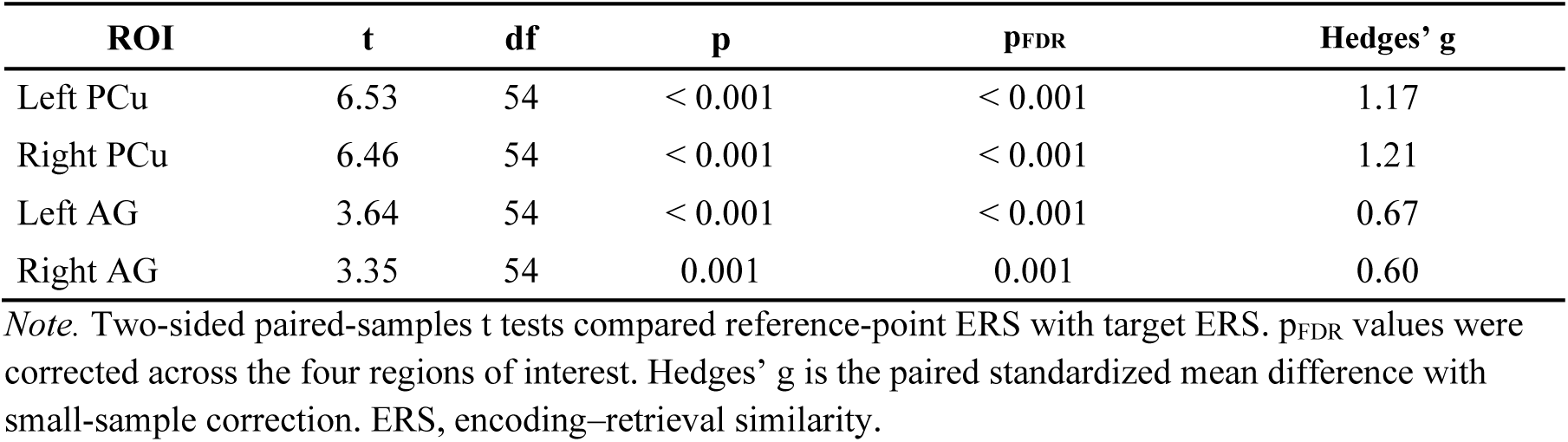
Reference-point ERS compared with target ERS.

**Supplementary Table 7.** Brain-region and hemisphere effects on reference-point and target ERS.

| ERS measure | F | dfn | dfd | p | p<sub>FDR</sub> | ηG<sup>2</sup> |
| --- | --- | --- | --- | --- | --- | --- |
| **Brain region** |  |  |  |  |  |  |
| Reference-point ERS | 5.50 | 1.00 | 54.00 | 0.023 | **0.046\*** | 0.014 |
| Target ERS | 1.59 | 1.00 | 54.00 | 0.213 | 0.213 | 0.006 |
| **Hemisphere** |  |  |  |  |  |  |
| Reference-point ERS | 0.31 | 1.00 | 54.00 | 0.578 | — | 0.001 |
| Target ERS | 0.08 | 1.00 | 54.00 | 0.774 | — | 0.000 |
| **Brain region × hemisphere** |  |  |  |  |  |  |
| Reference-point ERS | 0.49 | 1.00 | 54.00 | 0.488 | — | 0.001 |
| Target ERS | 1.54 | 1.00 | 54.00 | 0.220 | — | 0.002 |

*Note.* Separate $2 \times 2$ repeated-measures ANOVAs were conducted for reference-point ERS and target ERS, with brain region (precuneus versus angular gyrus) and hemisphere (left versus right) as within-participant factors. p<sub>FDR</sub> values were corrected across the two planned brain-region effects; significant FDR-corrected p values are shown in bold. dfn, numerator degrees of freedom; dfd, denominator degrees of freedom; ηG<sup>2</sup>, generalized eta squared; ERS, encoding–retrieval similarity.

**Supplementary Table 8.**
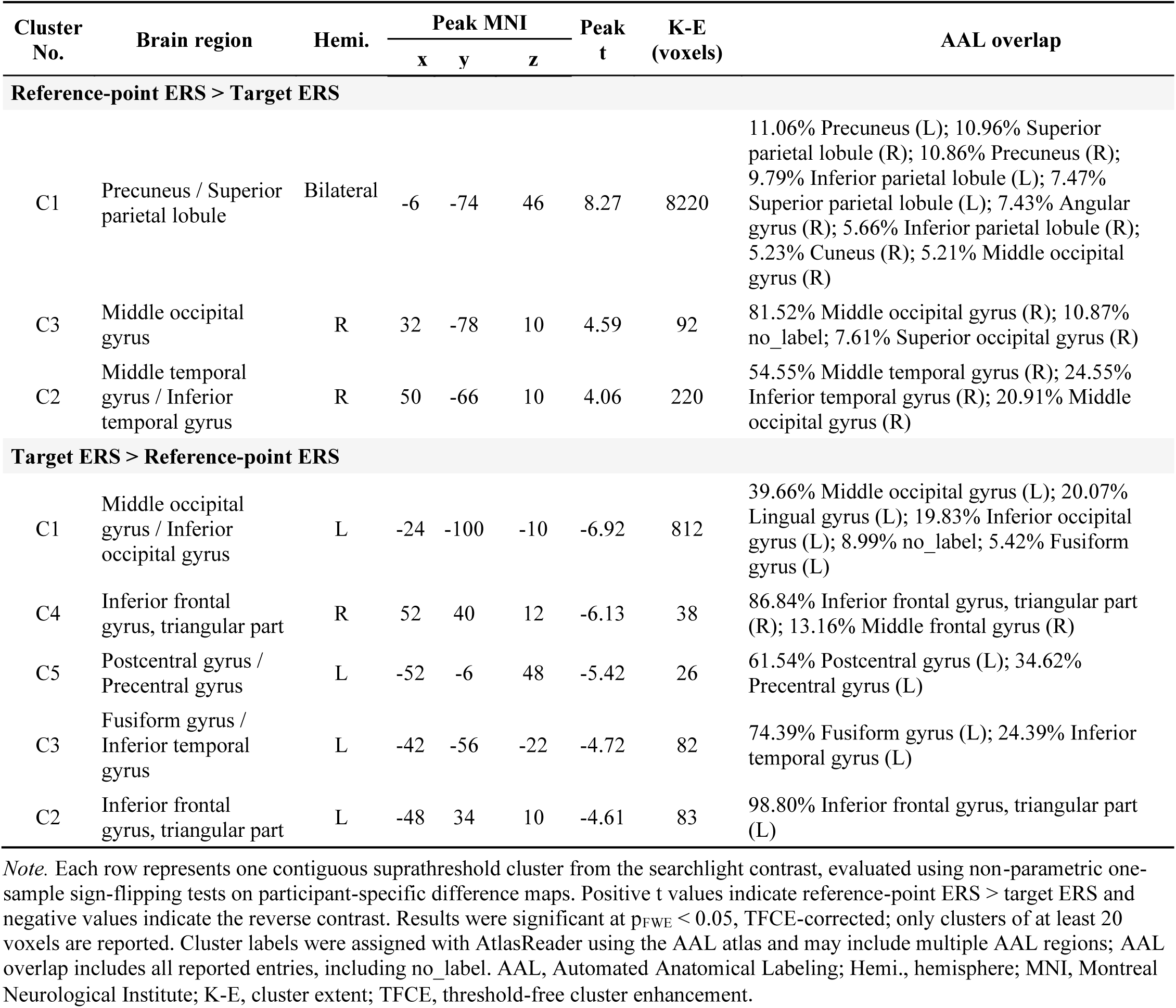
Searchlight clusters for the reference-point ERS versus target ERS contrast.

**Supplementary Table 9.** Reference-to-target path ERS predicting trial-wise temporal-order judgment correctness.

| ROI | Predictor | $\beta$ | z | Odds ratio | 95% CI | p | p<sub>FDR</sub> |
| --- | --- | --- | --- | --- | --- | --- | --- |
| Left PCu | reference-to-target path ERS | 0.20 | 2.70 | 1.22 | [1.06, 1.41] | 0.007 | **0.007\*\*** |
| Right PCu | reference-to-target path ERS | 0.25 | 3.31 | 1.28 | [1.11, 1.48] | < 0.001 | **0.004\*\*** |
| Left AG | reference-to-target path ERS | 0.20 | 2.69 | 1.22 | [1.06, 1.41] | 0.007 | **0.007\*\*** |
| Right AG | reference-to-target path ERS | 0.21 | 2.87 | 1.23 | [1.07, 1.42] | 0.004 | **0.007\*\*** |

*Notes for Supplementary Tables 9–17.* Values are standardized fixed-effect estimates ( $\beta$ ) from region-of-interest mixed-effects logistic regression models of trial-wise temporal-order judgment correctness. Each model included trial type as a categorical fixed effect and participant as a random intercept. Wald z and p values test the named fixed effect; odds ratios are $\exp(\beta)$ . p<sub>FDR</sub> values were corrected across the four predefined regions of interest separately for each predictor. Symbols denote FDR-corrected significance: ~p<sub>FDR</sub> < 0.10, \*p<sub>FDR</sub> < 0.05, \*\*p<sub>FDR</sub> < 0.01, \*\*\*p<sub>FDR</sub> < 0.001. CI, confidence interval; ERS, encoding–retrieval similarity.

**Supplementary Table 10.**
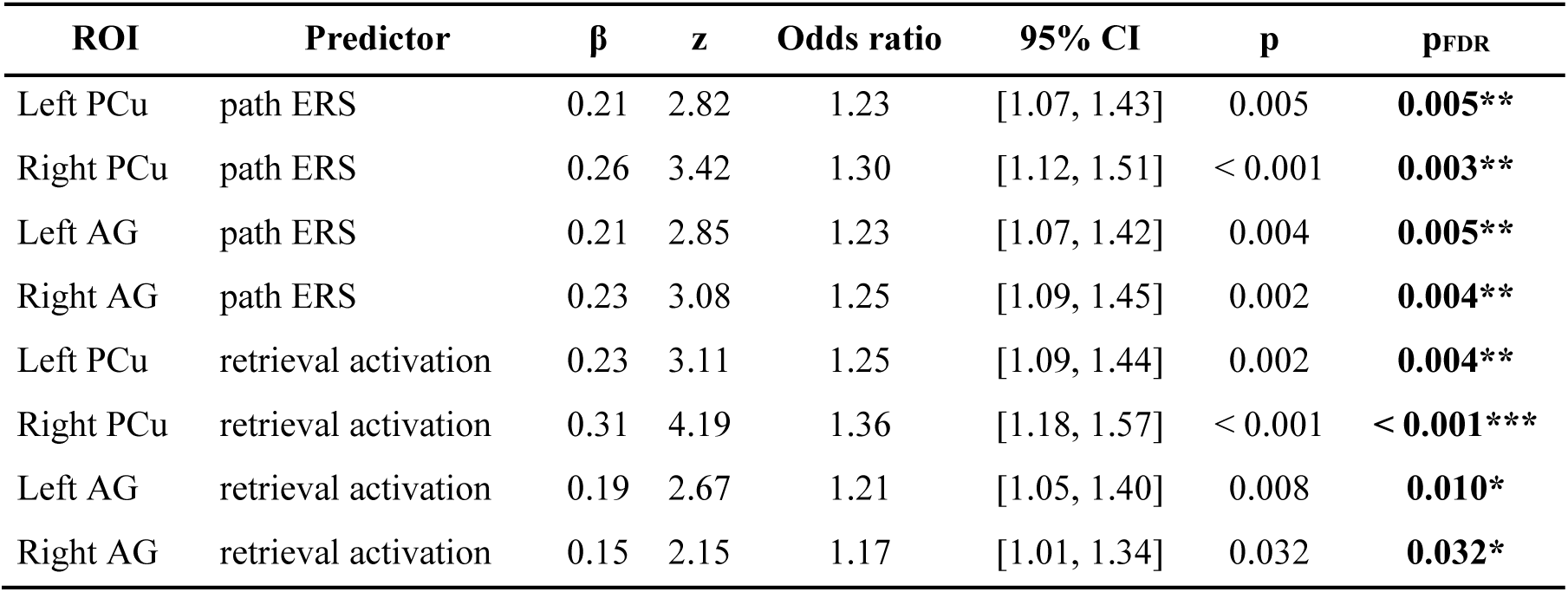
Path ERS predicting trial-wise temporal-order judgment correctness after adjustment for retrieval activation.

**Supplementary Table 11.** Reference-point ERS predicting trial-wise temporal-order judgment correctness.

| ROI | Predictor | $\beta$ | z | Odds ratio | 95% CI | p | p<sub>FDR</sub> |
| --- | --- | --- | --- | --- | --- | --- | --- |
| Left PCu | reference-point ERS | 0.09 | 1.25 | 1.09 | [0.95, 1.26] | 0.213 | 0.213 |
| Right PCu | reference-point ERS | 0.14 | 1.90 | 1.15 | [1.00, 1.32] | 0.057 | 0.116 |
| Left AG | reference-point ERS | 0.14 | 1.90 | 1.15 | [1.00, 1.32] | 0.058 | 0.116 |
| Right AG | reference-point ERS | 0.11 | 1.62 | 1.12 | [0.98, 1.29] | 0.105 | 0.140 |

**Supplementary Table 12.**
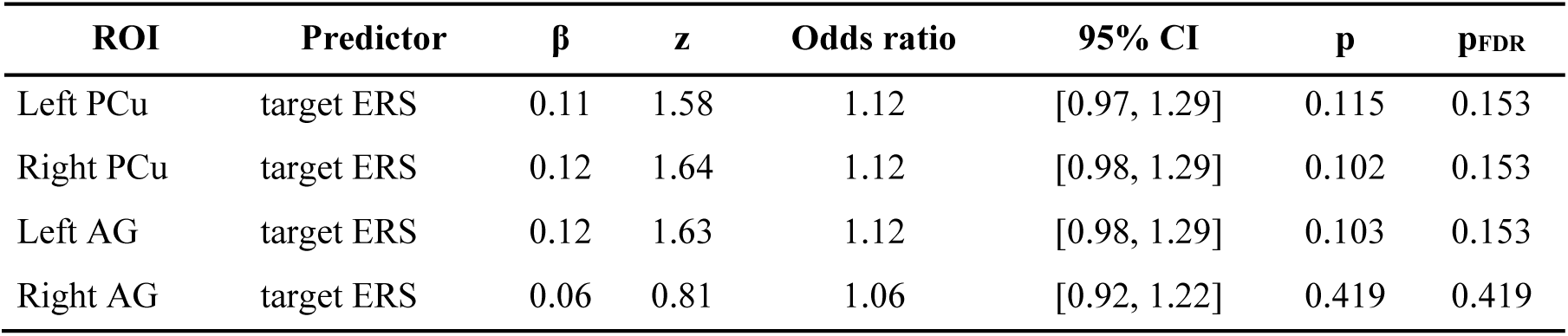
Target ERS predicting trial-wise temporal-order judgment correctness.

**Supplementary Table 13.** Reference-excluded path ERS predicting trial-wise temporal-order judgment correctness.

| ROI | Predictor | $\beta$ | z | Odds ratio | 95% CI | p | pFDR |
| --- | --- | --- | --- | --- | --- | --- | --- |
| Left PCu | reference-excluded path ERS | 0.16 | 2.17 | 1.17 | [1.02, 1.35] | 0.030 | **0.061**~ |
| Right PCu | reference-excluded path ERS | 0.17 | 2.30 | 1.18 | [1.03, 1.36] | 0.021 | **0.061**~ |
| Left AG | reference-excluded path ERS | 0.12 | 1.70 | 1.13 | [0.98, 1.30] | 0.089 | **0.089**~ |
| Right AG | reference-excluded path ERS | 0.14 | 1.97 | 1.15 | [1.00, 1.33] | 0.049 | **0.065**~ |

**Supplementary Table 14.**
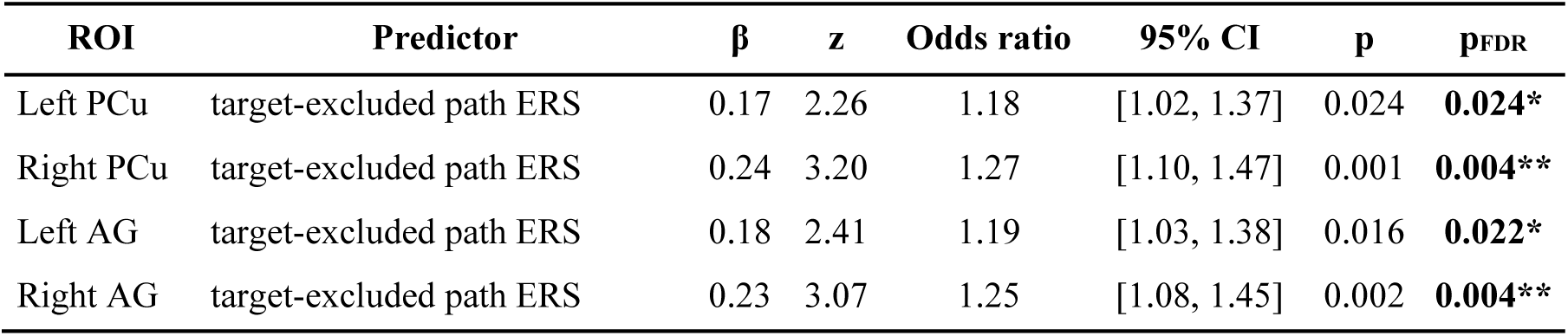
Target-excluded path ERS predicting trial-wise temporal-order judgment correctness.

| ROI | Predictor | $\beta$ | z | Odds ratio | 95% CI | p | pFDR |
| --- | --- | --- | --- | --- | --- | --- | --- |
| Left PCu | target-excluded path ERS | 0.17 | 2.26 | 1.18 | [1.02, 1.37] | 0.024 | **0.024** * |
| Right PCu | target-excluded path ERS | 0.24 | 3.20 | 1.27 | [1.10, 1.47] | 0.001 | **0.004** ** |
| Left AG | target-excluded path ERS | 0.18 | 2.41 | 1.19 | [1.03, 1.38] | 0.016 | **0.022** * |
| Right AG | target-excluded path ERS | 0.23 | 3.07 | 1.25 | [1.08, 1.45] | 0.002 | **0.004** ** |

**Supplementary Table 15.** Out-path ERS predicting trial-wise temporal-order judgment correctness.

| ROI | Predictor | $\beta$ | z | Odds ratio | 95% CI | p | pFDR |
| --- | --- | --- | --- | --- | --- | --- | --- |
| Left PCu | out-path ERS | 0.01 | 0.15 | 1.01 | [0.88, 1.16] | 0.881 | 0.991 |
| Right PCu | out-path ERS | 0.09 | 1.26 | 1.09 | [0.95, 1.26] | 0.209 | 0.836 |
| Left AG | out-path ERS | 0.00 | 0.01 | 1.00 | [0.87, 1.15] | 0.991 | 0.991 |
| Right AG | out-path ERS | 0.03 | 0.48 | 1.03 | [0.90, 1.19] | 0.629 | 0.991 |

**Supplementary Table 16.** Joint model of path ERS and out-path ERS predicting trial-wise temporal-order judgment correctness.

| ROI | Predictor | $\beta$ | z | Odds ratio | 95% CI | p | pFDR |
| --- | --- | --- | --- | --- | --- | --- | --- |
| Left PCu | path ERS | 0.20 | 2.70 | 1.22 | [1.06, 1.41] | 0.007 | **0.007\*\*** |
| Right PCu | path ERS | 0.24 | 3.25 | 1.27 | [1.10, 1.48] | 0.001 | **0.005\*\*** |
| Left AG | path ERS | 0.20 | 2.70 | 1.22 | [1.06, 1.41] | 0.007 | **0.007\*\*** |
| Right AG | path ERS | 0.21 | 2.84 | 1.23 | [1.07, 1.42] | 0.004 | **0.007\*\*** |
| Left PCu | out-path ERS | 0.01 | 0.19 | 1.01 | [0.88, 1.17] | 0.846 | 0.858 |
| Right PCu | out-path ERS | 0.08 | 1.11 | 1.08 | [0.94, 1.24] | 0.268 | 0.858 |
| Left AG | out-path ERS | -0.01 | -0.18 | 0.99 | [0.86, 1.14] | 0.858 | 0.858 |
| Right AG | out-path ERS | 0.02 | 0.29 | 1.02 | [0.89, 1.17] | 0.774 | 0.858 |

**Supplementary Table 17.** Joint model of path ERS, out-path ERS and retrieval activation predicting trial-wise temporal-order judgment correctness.

| ROI | Predictor | $\beta$ | z | Odds ratio | 95% CI | p | pFDR |
| --- | --- | --- | --- | --- | --- | --- | --- |
| Left PCu | path ERS | 0.21 | 2.84 | 1.24 | [1.07, 1.43] | 0.005 | **0.005\*\*** |
| Right PCu | path ERS | 0.26 | 3.34 | 1.29 | [1.11, 1.50] | < 0.001 | **0.003\*\*** |
| Left AG | path ERS | 0.21 | 2.84 | 1.23 | [1.07, 1.42] | 0.005 | **0.005\*\*** |
| Right AG | path ERS | 0.22 | 3.04 | 1.25 | [1.08, 1.44] | 0.002 | **0.005\*\*** |
| Left PCu | out-path ERS | 0.05 | 0.69 | 1.05 | [0.91, 1.21] | 0.493 | 0.712 |
| Right PCu | out-path ERS | 0.13 | 1.75 | 1.14 | [0.98, 1.32] | 0.081 | 0.322 |
| Left AG | out-path ERS | 0.01 | 0.12 | 1.01 | [0.88, 1.16] | 0.904 | 0.904 |
| Right AG | out-path ERS | 0.04 | 0.62 | 1.05 | [0.91, 1.20] | 0.534 | 0.712 |
| Left PCu | retrieval activation | 0.23 | 3.18 | 1.26 | [1.09, 1.46] | 0.001 | **0.003\*\*** |
| Right PCu | retrieval activation | 0.33 | 4.39 | 1.39 | [1.20, 1.60] | < 0.001 | **< 0.001\*\*\*** |
| Left AG | retrieval activation | 0.19 | 2.67 | 1.21 | [1.05, 1.40] | 0.008 | **0.010\*** |
| Right AG | retrieval activation | 0.16 | 2.22 | 1.17 | [1.02, 1.35] | 0.027 | **0.027\*** |

